# Spatial and Temporal Coordination of Force-generating Actin-based Modules Drives Membrane Remodeling *In Vivo*

**DOI:** 10.1101/2023.12.04.569944

**Authors:** Marco Heydecker, Akiko Shitara, Desu Chen, Duy Tran, Andrius Masedunskas, Muhibullah Tora, Seham Ebrahim, Mark A. Appaduray, Jorge Luis Galeano Niño, Abhishek Bhardwaj, Kedar Narayan, Edna C. Hardeman, Peter W. Gunning, Roberto Weigert

**Affiliations:** Laboratory of Cellular and Molecular Biology, Center for Cancer Research, National Cancer Institute, National Institutes of Health, Bethesda, MD, USA; School of Biomedical Sciences, University of New South Wales Sydney, Sydney, NSW, Australia; Department of Molecular Physiology and Biological Physics, Center for Membrane and Cell Physiology, University of Virginia School of Medicine, Charlottesville, VA, USA; Center for Molecular Microscopy, Center for Cancer Research, National Cancer Institute, National Institutes of Health, Bethesda, MD, USA and Cancer Research Technology Program, Frederick National Laboratory for Cancer Research, Frederick, MD, USA; Department of Pharmacology, Asahi University School of Dentistry, 1851-1 Hozumi, Mizuho, Gifu 501-0296, Japan; NIDCR Imaging Core, National Institute of Dental and Craniofacial Research, National Institutes of Health, Bethesda, MD, USA; EMBL Australia, Single Molecule Science node, University of New South Wales Sydney, NSW, Australia

**Keywords:** Membrane remodeling, Actin, Intravital microscopy, Exocytosis

## Abstract

Membrane remodeling drives a broad spectrum of cellular functions, and it is regulated through mechanical forces exerted on the membrane by cytoplasmic complexes. Here, we investigate how actin filaments dynamically tune their structure to control the active transfer of membranes between cellular compartments with distinct compositions and biophysical properties. Using intravital subcellular microscopy in live rodents we show that: a lattice composed of linear filaments stabilizes the granule membrane after fusion with the plasma membrane; and a network of branched filaments linked to the membranes by Ezrin, a regulator of membrane tension, initiates and drives to completion the integration step. Our results highlight how the actin cytoskeleton tunes its structure to adapt to dynamic changes in the biophysical properties of membranes.

## Introduction

Cells continuously remodel their membranes to drive a broad variety of essential cellular events ^1–3^. Membrane remodeling requires the exquisite coordination of different processes that include: i) modification of the lipid bilayer composition ^4,5^, ii) interaction between membranes and proteins harboring curvature-generating domains (e.g. BAR-domains) ^6,7^, and iii) application of mechanical forces by cytoplasmic complexes including the cytoskeleton ^8–11^. In particular, the actomyosin cytoskeleton has been implicated in membrane remodeling during processes ranging from cell division and cell migration to membrane trafficking ^12–16^. Its ability to reshape membranes with different biophysical properties (e.g., plasma membrane domains or vesicular structures) is linked to the flexibility of actin to form structurally and functionally diverse modules, such as branched and linear filaments ^17,18^, and to engage with the various members of the myosin family of motor proteins ^19,20^. F-actin and nonmuscle myosin II (NMII) are perfect examples of this modularity, as they assemble into a range of force-generating structures including contractile rings, sarcomeres, arcs and lattices ^21–24^.

How the activation and assembly of actomyosin complexes is transduced into the deformation of membranes has been extensively investigated in *in vitro* model systems (i.e., cell-free systems, cell culture in 2D and 3D, organoids) but few studies have been conducted *in vivo*. Although reductionist systems allow ready manipulation of experimental conditions, they do not take into account the crucial roles played by the 3D tissue organization and unique cues coming from the vasculature, nervous system, and adjacent cells during the remodeling. The development of intravital subcellular microscopy (ISMic) has opened the door to the visualization of the dynamics of subcellular processes in live animals at a resolution comparable to that achieved in cell culture ^25–29^. This approach has enabled the dissection of mechanisms controlling membrane remodeling in the context of organ physiology ^30,31^, cancer ^32,33^, cell motility ^34,35^, and membrane trafficking ^26,36–38^. We used ISMic to investigate the role of the actomyosin cytoskeleton in the gradual integration of large exocytic vesicles (termed granules) into the apical plasma membrane (APM) of acinar cells in salivary glands and the pancreas during regulated exocytosis in vivo. This process requires the assembly of a novel actomyosin-based structure composed of two intertwined lattices formed by F-actin and NMII ^21,36,39^. We have established that the role of the NMII lattice during granule integration is linked to NMII contractile activity ^21,39^. On the other hand, although we determined that the F-actin lattice is essential for the process, its specific function is not fully understood beyond its contribution to myosin-driven contraction.

Here, we show that immediately after fusion with the APM, the granules become coated with a lattice that is formed by linear actin filaments that are nucleated by mDia1, a member of the formin family of actin nucleators ^40,41^, that co-assemble with Tpm3.1, a member of the tropomyosin family of actin-associated proteins ^42^. This step is followed by the assembly of a wave of branched filaments controlled by the Arp2/3 complex ^17,43^, which drives granule integration. Impairment of the lattice assembly through pharmacological inhibition or genetic ablation of mDia1 results in a significant increase in the diameter of the fused granules due to an uncontrolled flow of membranes from the APM and also fusion with adjacent granules (i.e., compound exocytosis ^44^). In contrast, impairment of the Arp2/3 complex or Ezrin, an actin-membrane linker and tension regulator in the ERM family (Ezrin, Radixin, Moesin) ^45–47^, inhibits formation of the wave and significantly delays membrane integration, without affecting the assembly of the F-actin lattice.

Based on our results, we propose a novel mechanism for the remodeling of membranes that is based on the sequential assembly of two distinct F-actin-based modules: a lattice formed by linear actin filaments, that prevents the addition of membranes to the fused granules; and a wave of branched actin filaments, that actively generates forces controlling the integration of the membranes.

## Results

### Morphologically distinct F-actin modules assemble during membrane integration

To investigate membrane remodeling and F-actin dynamics during regulated exocytosis we used ISMic imaging of GFP-LF/mTom mice, that express the F-actin probe GFP-LifeAct (GFP-LF) ^48^ and the membrane-targeted peptide mTomato (mTom) ^49^ (**Fig. 1A**). As previously shown, the stimulation of exocytosis using isoproterenol (ISOP) leads to the fusion of the secretory granules with the APM ^36^ (**Fig. 1A,B** arrows). The diffusion of mTom from the apical canaliculi into the membranes of the secretory granules and the recruitment of GFP-LF make it possible to follow the dynamics of F-actin during the integration process (**Fig. 1A**,**C**) (please note that the terms GFP-LF and F-actin will be used interchangeably) ^21,36,39^. F-actin was detected on the membranes 1-2 sec after the appearance of mTom (**Fig. 1C**) as previously reported ^36^. The diameter of the fused granules did not significantly decrease for the first 6 sec (see Methods, N=55 granules in 5 animals) (**Fig. 1C,D; Movie S1**) and during this time F-actin assembled into the previously described lattice ^21^ (**Fig. 1E** arrow). Over the following 10-15 sec the granules decreased in size with a calculated rate of reduction of the surface area of 0.37±0.02 µm^2^/sec (see Methods, N=55 granules in 5 animals). Notably, as the diameter of the granules began to decrease, the external diameter of the lattice did not significantly change. However, polymerization of actin continued and manifested as an increase in the thickness of F-actin around the granules (hereafter referred as the wave) (**Fig. 1C** arrows). In addition, we observed the assembly of a denser layer of F-actin at the interface with the granule membranes (**Fig. 1C** double arrowheads). High resolution images acquired in fixed salivary glands confirmed the existence of the dense GFP-LF structure that lies beneath the external lattice and that interfaces with the granule membrane (**Fig. 1F** arrowheads) (hereafter referred as the inner ring). Similar results were obtained in fixed glands excised from mTom mice and stained with phalloidin, thus ruling out any possible artifacts due to the use of the GFP-LF probe (**Fig. 1E,F**). Quantitative analysis confirmed that the F-actin thickness inversely correlated with the granule diameter as measured both in GFP-LF/mTom mice imaged by ISMic and in fixed glands excised from mTom mice (**Fig. 1G,H**). At around 20 sec after fusion, when the diameter of the granules reached 20% of the initial size, the external diameter of the F-actin lattice began to decline steadily until 50-60 sec when F-actin was no longer detectable (**Fig. 1C**,**D**). Due to the limitations in the resolution of our instrumentation we could not perform any reliable dynamic measurement of the granule diameter below 200 nm or visualize the process of F-actin disassembly ^21^. Our analysis of the integration was limited, therefore, to a range of diameters between 1.2 and 0.3 µm. Focus ion beam scanning electron microscopy (FIB-SEM) ^50^ confirmed that during the integration process the granules maintained roundness and positive curvature (**Fig. S1A-C** and see Discussion). Moreover, we observed small membranous profiles bulging from the APM (<300 nm) although we could not distinguish whether they were secretory or endocytic in nature (**Fig. S1A**). Finally, we determined that the distribution of the diameters of the fused granules agreed with the measurements performed using light microscopy (**Fig. S1D**).

**Figure 1.**
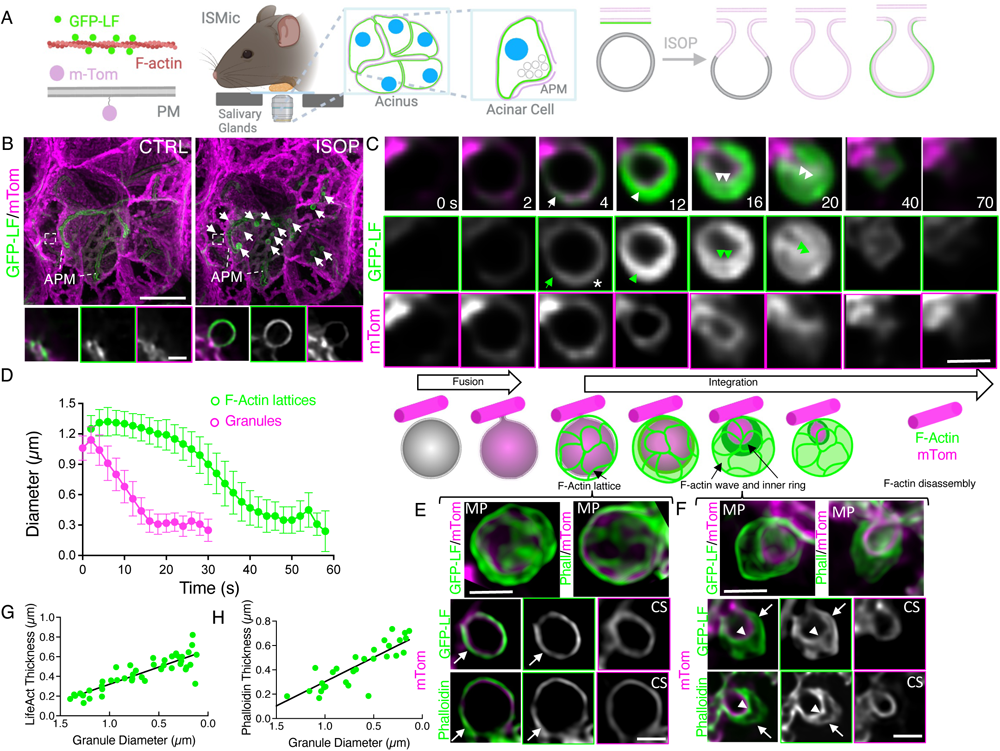
Dynamics of actin assembly on secretory granules during regulated exocytosis indicates multiple populations of F-actin. **A**. Diagram of the GFP-LF/mTom mouse and ISMic set up for the imaging of the submandibular salivary glands in live mice. Upon ISOP stimulation secretory granules fuse with the APM allowing the mTom (pink) to diffuse into the granule and the GFP-LF (green) to be recruited onto the membranes. GFP-LF/mTom (**B-F**) or mTom (**E,F,H**) mice were injected with 0.01 mg/kg ISOP. Acinar cells were either exposed and imaged by time-lapse spinning disk ISMic (**B-D**) or excised, fixed and processed for imaging using spinning disk microscopy (**E-H**). **B**. Acinar cells before (CTRL) and after stimulation (ISOP). Arrows in the right panel point to fused granules. mTom (magenta) and GFP-LF (green). Insets show a close-up of the APM and a fused granule. Bar 10 µm. **C**. Still images from a time-lapse of a secretory granule integration event (**Movie S1**). The focal plane was chosen to show the mid-cross section of the granule. Asterisk marks the first time point in which mTom is detected, arrows point to F-actin lattice, arrowheads to the F-actin thickening, and double arrowheads to the inner actin structure. Bar 1 µm. **D.** Lattice (green symbols) and granule (magenta symbols) diameters were measured. Data are averages +/-S.D. of 55 granules in 5 animals. **E,F.** High resolution spinning disk images of fused secretory granules before (**E**) and during (**F**) integration. Upper panels show maximal projections (MP) of the granules. Lower panels show cross sections (CS) of the granules. The lattice and the thicker layer of actin are highlighted by arrows and arrowheads, respectively. Bar 1 µm. **G,H.** Correlation between granule diameter and either LF (**G**: R^2^=0.7565 p<0.0001; N=25 granules in 3 animals) or phalloidin (**H**: R^2^=0.7407 p<0.0001; N=28 granules in 3 animals) thickness was calculated.

### mDia1 and the ARP2/3 complex are recruited sequentially onto the secretory granules after fusion with the APM

Our data suggest that two distinct F-actin-based populations control membrane integration. To establish whether the lattice and the wave of F-actin are assembled via different mechanisms, we determined which actin nucleators are recruited on the secretory granules after fusion with the APM. We used qPCR (**Fig. S2A**) to determine their expression in freshly isolated acinar cells, and indirect immunofluorescence *in vivo* to validate their recruitment on the fused secretory granules (**Fig. 2A,B; Fig. S2B**). We found that one member of the formin family of linear filament nucleators, mDia1 (**Fig. 2A**) ^40,41^, and components of the actin branching machinery that include two subunits of the Arp2/3 complex (ARPC2 and Arp2, **Fig. 2B** and **Fig. S2B**, respectively) ^43^, cortactin and N-WASP (**Fig. S2B**) ^51^ were expressed in acinar cells and recruited to the secretory granules after fusion with the APM. Under basal conditions these molecules localized either at the APM or in the cytoplasm, but not on the unfused granules (**Fig. S2C**).

**Figure 2.**
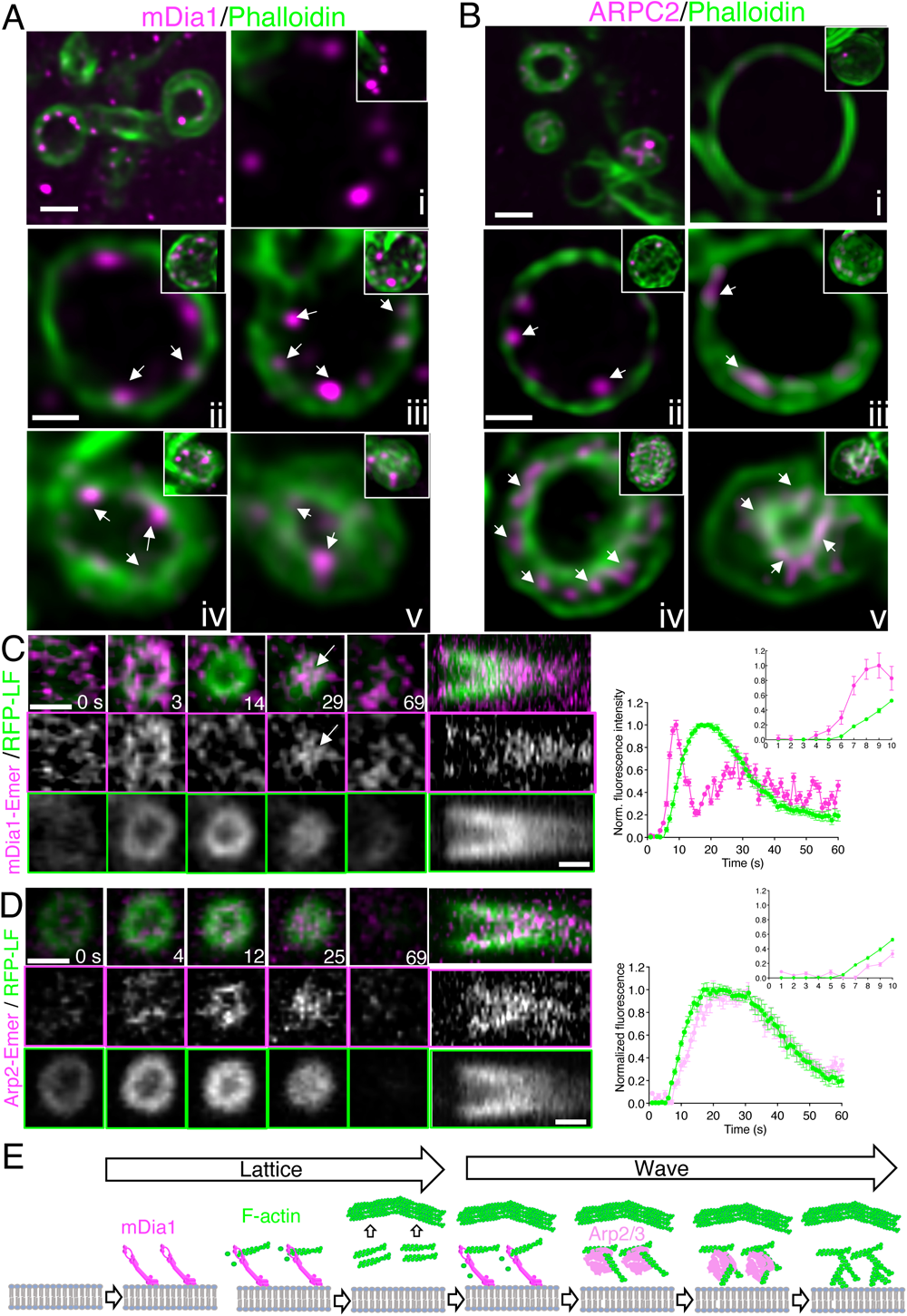
mDia1 and ARPC2 are recruited sequentially on fused secretory granules. **A,B.** WT mice were injected with 0.01 mg/kg ISOP and after 10 min salivary glands were processed for immunofluorescence and stained with Alexa488 phalloidin and antibodies against mDia1 (**A**) or ARPC2 (**B**). Images were acquired by spinning disk and processed. Cross sections of large granules before the wave (**i-ii**), intermediate size granules (**iii-iv**), and small granules (**v**). Insets show 3D view of the granules. Bar 1.5 µm (overview) and 0.5 µm (high magnification). **C**,**D.** Wistar rats were transfected through the salivary duct with RFP-LF and either mDia1-Emerald (**C**) or Arp2-Emerald (**D**). After 24 hr salivary glands were exposed, animals injected with 0.025 mg/kg ISOP and imaged using confocal ISMic. Still images from time-lapses (left panels; **Movie S2**), kymographs (center panels; **Movie S3**) and quantitation of recruitment of RFP-LifeAct, mDia1, and Arp2 (right panels). Arrow in **C** points to the second pool of mDia1 recruited on the granules. Data are averages +/- SD of 10 granules in 2 animals (mDia1/LifeAct) and 12 granules in 2 animals (Arp2/LifeAct). Bar 1 µm. **E**. Model of step-wise recruitment for linear and branched filaments.

mDia1 initially localized primarily in puncta associated with the secretory granules (**Fig. 2A**). The puncta partially overlapped with the F-actin lattice in larger granules (**Fig. 2Aii**), beneath the thick F-actin layer in intermediate size granules (**Fig. 2Aiii**), and with the inner F-actin ring in the smaller granules (**Fig. 2Aiv-v**). At all stages mDia1 preferentially localized at the interface with the membranes (**Fig 2Aii-v** arrows). Of note, we occasionally observed mDia1 puncta on granules that were devoid of phalloidin (**Fig. 2Ai**), suggesting that the recruitment of mDia1 precedes F-actin assembly. Similarly, we found that some large granules exhibiting the F-actin lattice were devoid of ARPC2, suggesting that the Arp2/3 complex is recruited and activated after the assembly of the lattice (**Fig. 2Bi**). Notably, in large granules ARPC2 localized beneath the lattice (**Fig. 2Bii** arrows) and as the thickening and the integration progressed APRC2 appeared to localize in the gap between the lattice and inner F-actin ring, and primarily associated with the latter (**Fig. 2Biii-v**).

These data are consistent with the initial recruitment of mDia1 on granule membranes immediately after fusion with the APM to initiate the assembly of linear actin filaments to form the lattice. This is followed by the recruitment of the Arp2/3 complex to initiate the F-actin wave composed of branched actin filaments, possibly in coordination with a second pool of mDia1 (**Fig. 2E**). To test this hypothesis, we used ISMic to determine the kinetics of actin nucleator recruitment on the granule membrane with respect to the assembly of F-actin. We co-transfected the acinar cells of the salivary glands of live rats with RFP-Lifeact (RFP-LF) and either mDia1 or Arp2 tagged with m-Emerald (**Fig. 2C,D**). Rats were chosen because: i) they can be transfected using an established procedure that doesn’t affect the structure/function of the acini ^52^, and ii) regulation of large granule exocytosis in salivary glands is similar to that of mice ^36^. We showed that both mDia1 and ARPC2 are present on secretory granules following ISOP stimulation in rats (**Fig. S2D**). Note, that due to the increased imaging depth in the rat salivary glands we used confocal rather than spinning disk microscopy with a partial loss in spatiotemporal resolution. As expected, both mDia1 and Arp2 were recruited to the fused granule membranes upon ISOP stimulation. mDia1 appeared on the granules ∼2-3 sec before the RFP-LF and its levels reached a peak after ∼10 sec (**Fig. 2C** left panels; **Movie S2**). This pool of transfected mDia1 localized primarily at the edge of the granules and overlapped with the F-actin lattice. We also observed a second pool of mDia1 that was recruited as the wave initiated inside the F-actin lattice (**Fig. 2C** left panels, arrow; **Movie S2).** On the other hand, Arp2 appeared on the membranes ∼5 sec after RFP-LF and its level peaked at around 20 sec, mirroring the thickening of F-actin (**Fig. 2D** left panels; **Movie S3**). Arp2 localized only in the thickened LF layer juxtaposed to the external lattice, confirming the findings obtained by immunofluorescence (**Fig. 2B**) and suggesting that the second pool of mDia1 could be required to nucleate linear filaments needed to initiate the branching actin network.

Importantly, our demonstration that two structurally distinct actin populations are sequentially assembled on the secretory granules was validated by the finding that only the F-actin forming on the external lattice co-localized with tropomyosin Tpm3.1 (**Fig. 3A-C; Movie S4)**. Tropomyosins form copolymers with linear actin filaments and we have shown that Tpm3.1 is recruited to the granule membrane with identical kinetics to F-actin during the first 5 sec of filament formation ^42,53^. Similarly, NMIIA was also recruited to the granule membrane surface, as shown using both ISMic and indirect immunofluorescence (**Fig. 3D-F, Movie S5**). As exocytosis progressed, Tpm3.1 and NMIIA remained enriched at the periphery and did not associate with the F-actin pool beneath the lattice, indicating that Tpm3.1 and NMIIA are not major components of the inner wave of F-actin (**Fig. 3B,D**).

**Figure 3.**
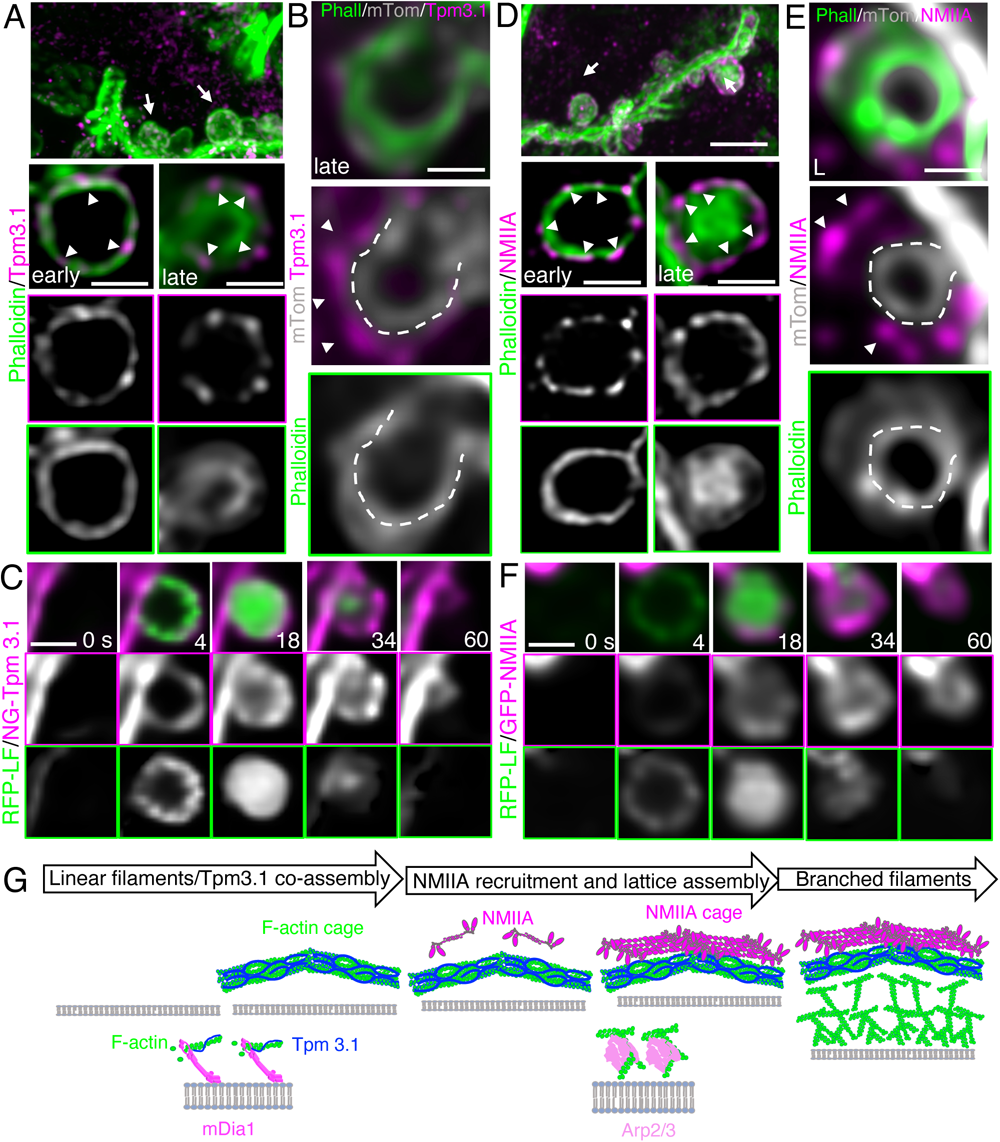
Tpm3.1 and NMIIA co-localize with the F-actin lattice, but not with the inner branched actin pool. WT (**A,D**), mTom (**B,E**), Tpm3.1-NG/RFP-LF (**C**), or RFP-LF/NMIIA (**F**) mice were injected with 0.01 mg/kg ISOP and salivary glands either exposed and imaged by ISMic (**C,F**) or after 10 min they were excised, processed for immunofluorescence, and stained with Alexa488 phalloidin and antibodies against Tpm3.1 (**A,B**) or NMIIA (**D,E**). **A,B & D,E.** Distribution of Tpm3.1 (**A,B** arrowheads), NMIIA (**D,E** arrowheads) on the secretory granules at early or late stage of integration. Dashed lines show the granules’ membranes. Bar 1 µm. **C,F.** Time-lapse of recruitment of Tpm3.1 and LifeAct (**C**; **Movie S4**) or NMIIA and LifeAct (**F**; **Movie S5**). Bar 1 µm. **G.** Model for assembly of Tpm3.1 and NMIIA on the external F-actin lattice.

### The mDia1-dependent F-actin lattice protects the fused granules from uncontrolled compound exocytosis

Our findings suggest that the fusion of the granule with the APM triggers the initial recruitment of mDia1 to the granule membrane and the assembly of actin/Tpm3.1 linear filaments, followed by the recruitment of the Arp2/3 complex and the generation of branched filaments (**Fig. 3G**). To determine the role of these two F-actin populations in the integration process, we used a Cre-lox approach to deplete the levels of either mDia1 or ARPC2, a key component of the Arp2/3 complex. Mice floxed for mDia1 (mDia1^fl/fl^) ^54^ or ARPC2 (ARPC2^fl/fl^) ^55^ were crossed with a strain expressing a tamoxifen (TMX)-inducible Cre recombinase under the control of the Mist1 promoter, which is expressed in salivary glands ^56^. These mice also harbor the mT/mGFP reporter to distinguish Cre-expressing cells (GFP positive) and to visualize the secretory granules after their fusion with the APM ^39,49^ (**Fig. S3A-C**; mDia1^fl/fl^ ^Mist^^1-Cre, mT/mG^, ARPC2^fl/fl^ ^Mist^^1-Cre, mT/mG^). Mice expressing the Mist1-Cre module and the mT/mGFP reporter (Ctrl^Mist^^1-Cre, mT/mG)^ were used as controls.

mDia1^fl/fl^ ^Mist1-Cre, mT/mG^ and Ctrl^Mist1-Cre, mT/mG^ mice were treated with TMX and imaged after 3-4 weeks. At this time the cellular levels of mDia1 were substantially reduced (**Fig. S3B**). Any attempt to further deplete mDia1 severely affected the endomembrane system of the acini (**Fig. S4A**). Under these conditions, in mDia1^-/-^ acinar cells, 37% +/- 7.1%. (102 granules in 8 animals) of the secretory granules began to expand immediately after ISOP-induced fusion with the APM and did not integrate for at least 90 sec (**Fig. 4A,B** left panels; **Fig. 4C**; **Movie S6**). Twenty percent (+/- 6.4%) of the granules underwent delayed integration; whereas, the remaining 43% integrated similarly to the controls (**Fig. 4C**). TMX administration or Cre expression did not induce this phenotype since the morphology and kinetics of integration of the secretory granules did not change in mDia1^+/+^ acinar cells in either mDia1^fl/fl^ ^Mist1-Cre, mT/mG^ (**Fig. S4B**) or Ctrl^Mist1-Cre, mT/mG^ (**Fig. 4A,B** right panels; **Fig. 4C**; **Movie S6**) mice treated with TMX. Moreover, mDia1 depletion did not result in defects in the assembly of F-actin on the APM or changes in its morphology (**Fig. S4C**), suggesting a direct impact of the linear F-actin associated with the granule on membrane integration. We investigated whether the heterogeneity in the integration phenotype in mDia1-/- cells was due to differences in the levels of F-actin assembly on the membrane of the granules, because of either variability in the extent of mDia1 depletion or compensation from other formins expressed in the acinar cells. We found that F-actin was consistently detected on granules with diameters below 1.5 µm (**Fig. S4D** lower panels); whereas, larger granules appeared to be either uncoated or partially coated with F-actin (**Fig. S4D** upper and center panels). We confirmed that mDia1 was not detected on the fused granules (**Fig. S3B**). To further substantiate these data and visualize the dynamics of F-actin recruitment when formin activity is impaired, we treated the GFP-LifeAct/mTom mice with SMIFH2, a drug that targets the FH2 domains shared by multiple members of the formin family ^57^. SMIFH2 treatment phenocopied the effects of mDia1 depletion on granule integration and F-actin assembly in mTom (**Fig. 4E,** N=103 granules in 8 animals), GFP-LifeAct/mTom (**Fig. 4F**), and WT strains (**Fig. S4E**). We observed that: i) F-actin was not recruited on the population of expanded granules that did not integrate (**Fig. 4F** upper panels), and ii) a partial recruitment of F-actin occurred at later time points on a subset of expanded granules, coinciding with the onset of the integration process (**Fig. 4F** center panels). Consistently, mDia1 and Arpc2 were recruited on these granules and most likely they were partially functional since we detected enlarged lattice-like structures (**Fig. 4G**) but did not observe any F-actin wave (**Fig. 4F** center panels). This finding suggests that a limited amount of active formins could still promote the assembly of the lattice but do not support the assembly of the Arp2/3-mediated branched filaments. However, under these conditions NMIIA localized on the F-actin-coated large granules (**Fig. 4G**) and possibly contributed to drive the integration, as previously reported ^48^. Finally, as expected, the remaining granules recruited F-actin, did not expand, and underwent normal integration (**Fig. 4F** lower panels).

**Figure 4.**
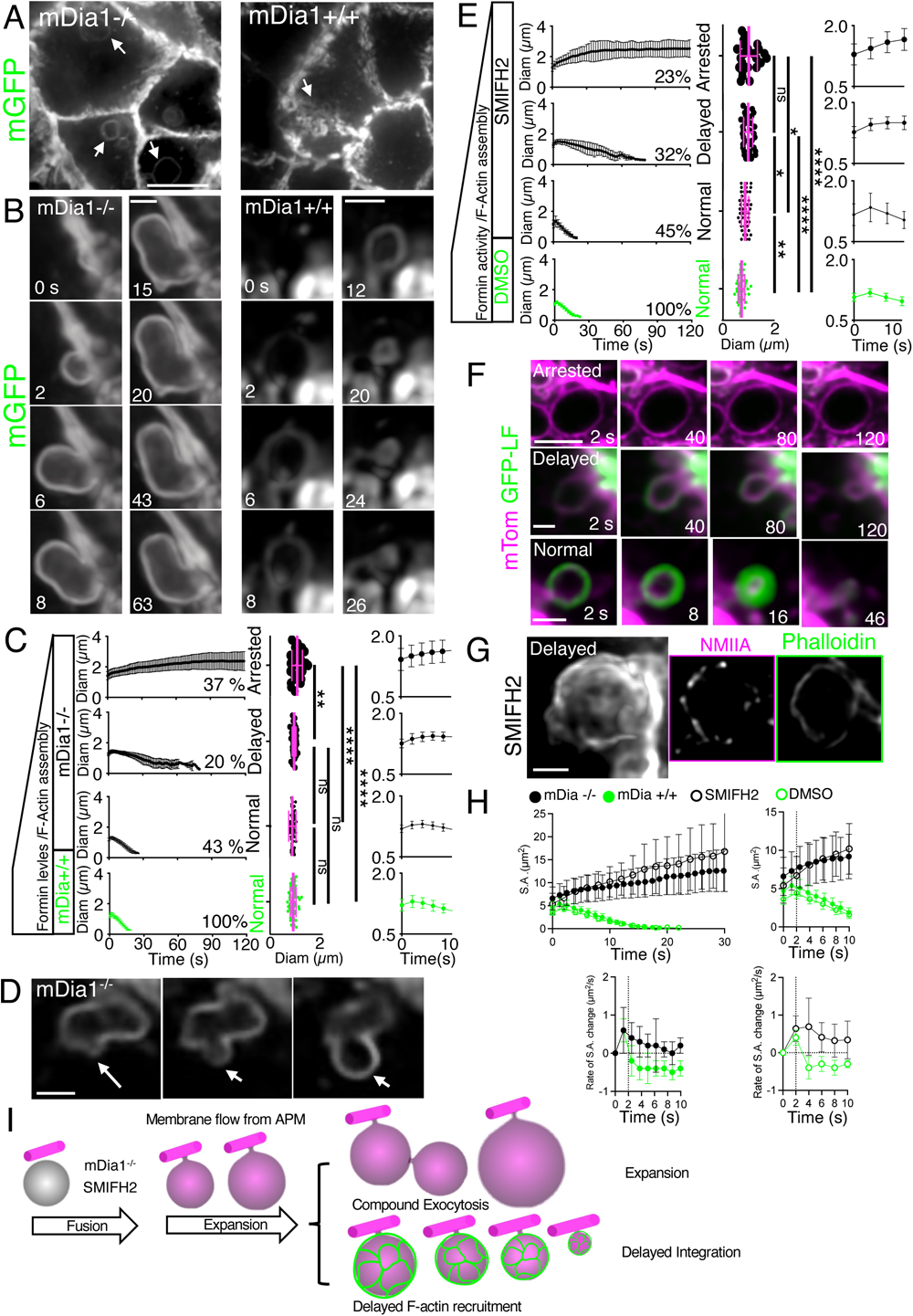
Depletion of mDia1 or pharmacological inhibition of formins affects the assembly of the F-actin lattice and the stabilization of the fused granules. mDia1^fl/fl^ ^Mist1-Cre, mT/mG^ (**A,B** left panels, **C,D,H**) or Ctrl^Mist^^1-Cre, mT/mG^ (**A,B** right panels, **C,H**) mice were treated with TMX. After 4 weeks salivary glands were exposed, mice injected with 0.01 mg/kg ISOP and imaged by ISMic (**A-D**, **H**). **A.** Expanded granules in mDia1-/-(arrows) and normal granule in mDia1+/+ (arrowheads) acinar cells. Bar 5 µm. **B.** ISMic time-lapse of granule integration showing expansion (left panels). Bar 1 µm. **C.** Time course (left panels) of granule diameter measured in mDia1-/- (black symbols; N=100 granules in 8 animals) and mDia1+/+ (green symbols; N=65 granules in 5 animals) cells. Diameters of secretory granules right after fusion (center panels). One-way ANOVA (****p<0.0001, ***p<0.001, **p<0.01, *p<0.1). Insets show diameter of the granules in the first 10 sec after fusion (right panels). Data are averages +/- S.D. **D**. Time-lapse of compound exocytosis. Bar 1 µm. **E-H.** mTom (**E,H**), GFP-LF/mTom (**F**) or WT (**G**) mice were treated with 200 µm SMIFH2 (**E-G**) or DMSO (vehicle, **E**), injected with 0.01 mg/kg ISOP, and salivary glands imaged by ISMic or fixed, processed for immunofluorescence, and imaged by spinning disk microscopy. **E**. Time course (left panels) of granule integration was measured in SMIFH2- (black symbols; N=103 granules in 8 animals) and DMSO-treated (vehicle, green symbols; N=48 granules in 5 animals) mice. Diameters of the secretory granules after fusion (center panels). One-way ANOVA (****p<0.0001, ***p<0.001, **p<0.01, *p<0.1). Insets show the diameter of the granules in the first 10 sec after fusion (right panels). Data are averages +/- S.D. **F**. Time-lapse of expanded granules in acinar cells. Upper panels - Expanded granule (magenta) that fails to recruit GFP-LF and does not integrate. Bar 3 µm. Center panels - Expanded granule that recruits GFP- LF (green) and undergoes delayed integration. Bar 1 µm. Lower panels - Granules that undergo normal integration. Bar 1 µm. **G.** Expanded granules undergoing delayed integration labeled with phalloidin (green) and an antibody against NMIIA (magenta). Left panel shows the F-actin lattice. Bar 1 µm. **H**. Surface area (S.A.) and rate of changes in S.A. were calculated for the mice described in **C** and **E**. **I**. Model for formin depletion/impairment.

Next, we addressed the mechanism responsible for the expansion of granules in the absence of the F-actin lattice immediately after their fusion with the APM. We envisioned two non-mutually exclusive mechanisms. The first is based on the fact that as the fusion pore forms, membrane is driven from the APM into the granule by a gradient of membrane tension, as previously shown during granule exocytosis in mast cells and in model membranes ^58,59^. Notably, in the first few seconds after fusion, the surface area of granules increased at a similar rate under both control conditions and when formin was impaired (**Fig. 4H**).

On the other hand, when F-actin recruitment was impaired the granule surface area steadily increased over time with absolute rates similar to those measured during the integration process (**Fig. 4H**). Although we cannot measure the overall surface area of the APM due to the high number of villi and folds present in its lumen ^60^, it is unlikely that a 3-4-fold increase in granule surface area (**Fig. 4H**) can be attributable solely to a flow of lipid from the APM. Indeed, we occasionally observed membranes being transferred from adjacent granules undergoing compound exocytosis ^44,61^ (**Fig. 4D**).

Together, these data support the hypothesis that the lattice formed by mDia1-dependent linear filaments provides a scaffold to stabilize the membrane of the granules after their fusion with the APM, and to prevent their expansion due to membrane transfer from either the APM or via compound exocytosis (**Fig. 4I**).

### The Arp2/3 complex controls the integration of secretory granules

We next addressed the role of the F-actin wave associated with the Arp2/3 complex and its integration with the function of the lattice. ARPC2^fl/fl^ ^Mist1-Cre, mT/mG^ mice were treated with TMX such that after 3-4 weeks the level of ARPC2 was below the level of detection (**Fig. S3C**). In ISOP-stimulated ARPC2^-/-^ acinar cells the granules fused with the APM and did not increase in size; however, their integration was significantly delayed (68 sec +/- 15 sec [N=46 granules in 4 animals] vs 16 sec +/- 1 sec [N=61 granules in 5 animals]) (**Fig. 5A-C**; **Movie S7**), and the diameter of the APM was not affected (**Fig. S4F**). Strikingly, the F-actin wave and inner ring were not observed although the structure of both the F-actin and NMIIA lattices were not affected (**Fig. 5D**). Accumulation of granules was occasionally observed in unstimulated ARPC2^-/-^ cells (**Fig. S4G**).

**Figure 5.**
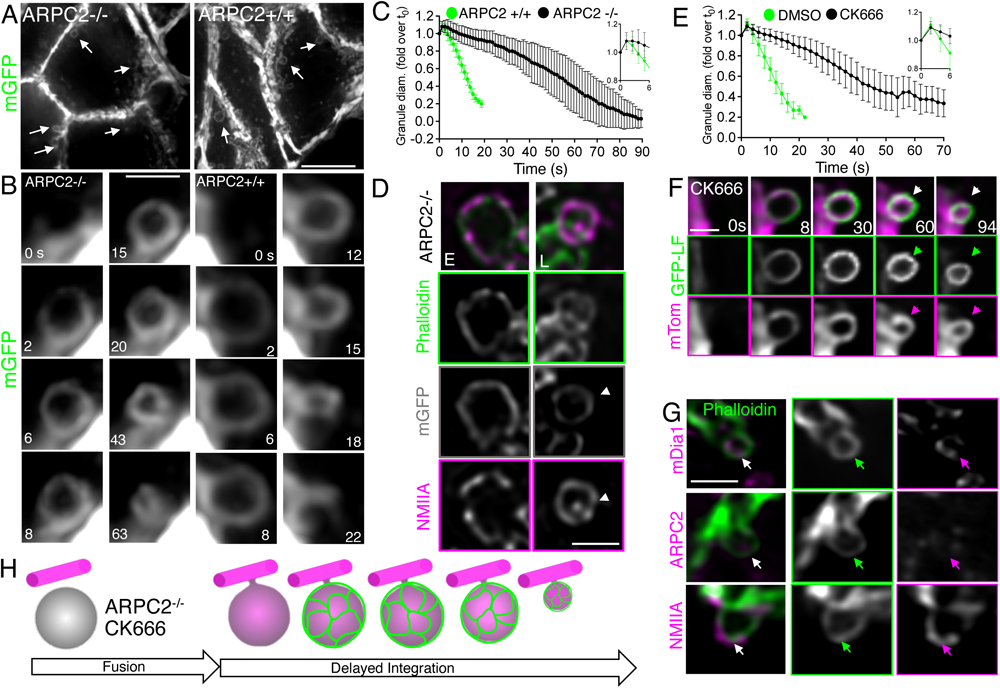
The Arp2/3 complex controls the wave of F-actin on the fused granules and their integration into the APM. **A-D.** ARPC2^fl/fl^ ^Mist1-Cre, mT/mG^ (**A,B** left panels, **C,D**) or Ctrl^Mist1-Cre, mT/mG^ (**A,B** right panels) were treated with TMX. After 4 weeks salivary glands were exposed, mice were injected with 0.01 mg/kg ISOP and imaged by ISMic (**A-C**) or processed for indirect immunofluorescence, labelled with Alexa-647 phalloidin, and antibodies directed against NMIIA, and imaged by spinning disk (**D**). **A.** Granules in ARPC2-/- and ARPC2+/+ acinar cells (arrows). Bar 5 µm. **B.** ISMic time-lapse of granule integration. Bar 1 µm. **C.** Time course of granule integration was measured in ARPC2-/- (green solid symbols) and ARPC2+/+ (green open symbols) cells. Data are averages +/- S.D. (ARPC2-/-: N=44 granules in 4 animals; ARPC2+/+: N=61 granules in 5 animals). **D**. Phalloidin and NMIIA staining of granules at early and later stage of integration. Bar 1 µm. **E-G**. GFP-LF/mTom (**E,F**) or WT (**G**) mice were treated with 200 µm CK666 or DMSO (vehicle) and injected with 0.01 mg/kg ISOP. Salivary glands were either imaged by ISMic (**E,F**) or processed for indirect immunofluorescence labelled with Alexa- 647 phalloidin and antibodies directed against mDia1, Arpc2, or NMIIA, and imaged by spinning disk (**G**). **E.** Time course of granule integration was measured for CK666- (green solid symbols) and vehicle-treated (DMSO, green open symbols) cells. Data are averages +/- S.D. (CK666: N=44 granules in 5 animals; DMSO: N=48 granules in 5 animals). Bar 1 µm. **H.** Model of the effect of Arp2/3 inhibition on granule integration.

The effects of the ARPC2 depletion were validated by acute inhibition of branched actin assembly following administration of the Arp2/3 inhibitor CK666 (**Fig. 5E-G**, ^62^). The integration of the granules was slowed without any effect on their initial diameter (**Fig. 5E**; **Movie S8**) and the F-actin thickening was largely abolished (**Fig. 5F**). Notably, consistent with this phenotype, the recruitment of ARPC2 was affected by CK666 treatment (**Fig. 5G**); whereas, the recruitment of mDia1 and NMIIA were not (**Fig. 5G**). These data suggest that the Arp2/3-complex-dependent branched network facilitates the integration process but is not essential (see Discussion). This raises the question: how does the branched actin filament network control the integration of the secretory granules?

### Ezrin is recruited on the secretory granules and crosslinks F-actin to the membranes to control integration

Actin filaments are known to exert forces on cellular membranes and control membrane tension through proteins such as the members of the ERM (Ezrin-Radixin-Moesin) family of linkers ^46^. We found that Ezrin and Radixin, which are expressed in epithelial cells ^63^, were localized at the APM, while Moesin exhibited a weak plasma membrane staining (**Fig. S5A**). However, only Ezrin and Radixin were recruited to fused secretory granules upon ISOP stimulation (**Fig. 6A**, **Fig. S5B**). In large granules Ezrin localized beneath the F-actin lattice and became associated primarily with the inner ring at the membrane interface as the integration progressed (**Fig. 6B**). On the other hand, Radixin localized only on the larger granules (**Fig. S5B**). Ezrin and Radixin are recruited to membranes via their FERM domains ^63^ and bind to F-actin via their C-terminal domains, which become available upon phosphorylation of conserved T567 and T654 residues, respectively ^64,65^. We could only confirm that Ezrin was phosphorylated on the fused secretory granules (**Fig. 6C,D**). To decouple F-actin from the membranes we used NSC 668394 (NSC), an inhibitor of Ezrin phosphorylation ^64^. Immunofluorescence confirmed that phosphorylation of Ezrin was significantly diminished in mice treated with NSC (**Fig. S5D,E**). Notably, treatment of GFP-LF/mTom mice with NSC did not inhibit the F-actin wave but prevented the assembly of the inner actin ring around the granules and delayed their integration with a reduced rate of surface area reduction (**Fig. 6E,F**; **Movie S9**). Under these conditions, Ezrin associated with the granules (**Fig. 6G**), both the F-actin and NMIIA lattices were not affected (**Fig. 6H** left panels), and there were no significant effects on the initial increase in surface area, suggesting that neither Ezrin or Radixin play a role at earlier time points (**Fig. 6F**). Finally, NSC did not affect the recruitment of the ARPC2 complex on the granules (**Fig. 6H** right panels). Overall, these findings suggest that the interaction between the branched filaments forming the inner ring and the membrane of the granules is mediated by Ezrin and are required to control granule integration (**Fig. 6I**).

**Figure 6.**
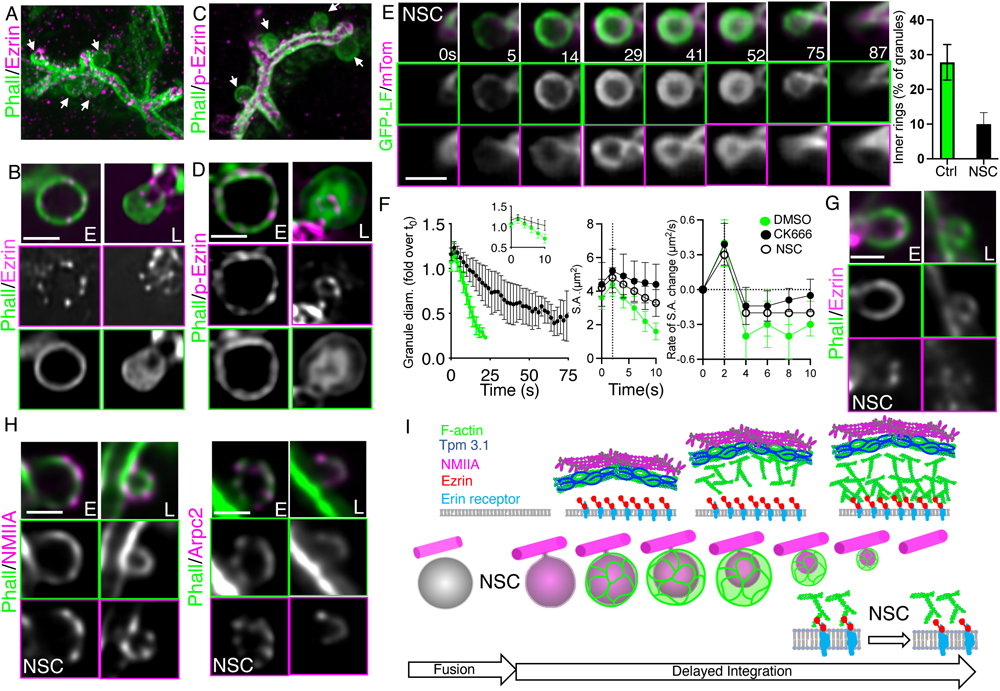
Inhibition of Ezrin delays the integration of the secretory granules. WT (**G,H**) or GFP-LF/mTom (**E,F**) mice were treated with 250 µM of NSC668394 or left untreated (**A-D**). Animals were stimulated with 0.01 mg/kg ISOP and then processed for immunofluorescence (**A-D,G,H**) or ISMic (**E,F**). **A-D.** Immunofluorescence staining of phalloidin (green), Ezrin (magenta, **A,B**) and p-Ezrin (magenta, **C,D**). **E.** ISMic time-lapse of granule integration (left panels) and quantification of % of granules exhibiting the F-actin inner ring (right panels). Bar 1 um. **F**. Time course of granule integration. Diameters, S.A. and rate of S.A. changes were measured. Data are averages +/- SD (NSC: N=36 granules in 4 animals; CK666: N=44 granules in 5 animals in Fig. 5); DMSO: N=48 granules in 4 animals). **G,H.** Immunofluorescence staining of phalloidin (green) and Ezrin (magenta, **G**), NMIIA (magenta, **H** left panel), or ARPC2 (magenta, **H** right panel). **I.** Model for Ezrin’s role during granule integration.

## Discussion

We propose a mechanism for the remodeling of membranes during exocytosis in living animals that is regulated by the coordinated action of two distinct force-generating F-actin-based modules. The first module is initiated by the rapid recruitment of the formin mDia1 on discrete foci of the membranes of the fused granules. mDia1 nucleates a population of linear actin filaments that co-polymerizes with Tpm3.1 ^66^ and assembles around the granules into lattice-like structures (**Fig. 7**). This lattice is intimately associated with a NMII lattice (**Fig. 3**, ^21^). The second module is initiated by the recruitment of the Arp2/3 complex that assembles a population of filaments composed of branched actin that interact with the membranes via Ezrin, a member of the ERM family of actin-membrane linkers ^67,68^ (**Fig. 7**). We predict that the F-actin lattice cannot serve as a seed to initiate branched filaments, since tropomyosins inhibit the ability of the Arp2/3 machinery to bind to actin ^69^. Although Arp2/3 has been shown to *de novo* initiate actin filaments ^70^, it is also possible that the second pool of mDia1 recruited on the membranes beneath the lattice provides the template mother filaments for the Arp2/3 complex (**Fig. 2**).

**Figure 7.**
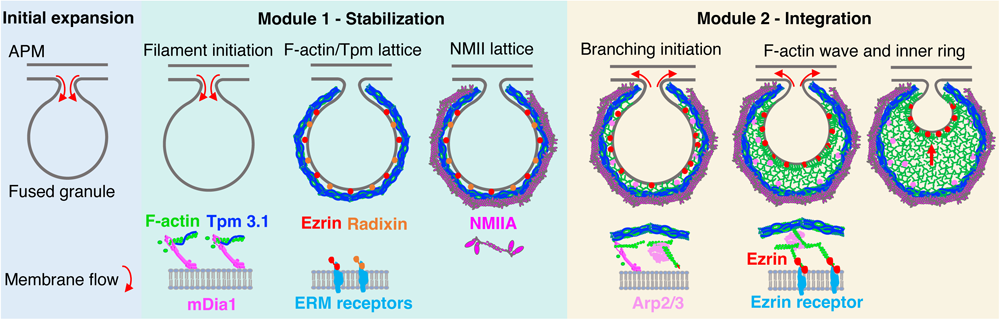
Model for the coordinated action of the two F-actin modules to control the integration of fused secretory granules. After fusion the secretory granules undergo an expansion most likely due to a flow of membrane from the APM. The recruitment of the first actin-based module (composed of a mDia1-dependent F-actin/Tpm3.1 lattice and a NMMIA lattice) stabilizes the membranes counteracting the flow. This step is followed by the assembly of the second module composed of Arp2/3-dependent branched filaments that interact with the ERM protein Ezrin, which triggers the integration of the granules’ membranes.

How do these modules control the remodeling of the membranes of the granules after their fusion with the APM? We envision that *in vivo* the integration step is driven by the progressive transfer of membranes from the secretory granules to the APM. Impairment of mDia1, which inhibits the early assembly of the F-actin/Tpm3.1 lattice without affecting the NMIIA lattice, results in the expansion of the granules right after fusion. This finding is consistent with both theoretical models and experimental data in mast cells where exocytic vesicles first transiently fuse with the plasma membrane to release quanta of cargo molecules and next are released into the cytoplasm (kiss and run). During this process, which does not involve F-actin coating of the fused granules, membranes are transferred from the plasma membrane to the exocytic vesicles through convective flow via the fusion pore ^58,59^. Therefore, we propose that the primary role of the F-actin lattice is to prevent the influx of membrane from the APM (**Fig. 4**). This function could require the lattice being linked to the membrane by Ezrin and/or Radixin, at least at the very early stages, although the NSC treatment did not induce any expansion of the granules and it did not affect the lattice possibly due to the assembly of the NMIIA lattice. This is consistent with the fact that: i) the F-actin lattice is still preserved when physically dissociated from the granule membrane at later stage of integration, and ii) ablation of NMII or inhibition of its contractile activity also induce an expansion of the granules although to a smaller extent ^21,39^. Moreover, in the absence of F-actin we observed a 2-to 3-fold increase in the granule diameter at later time points, suggesting that this large excess of membrane is provided by fusion with adjacent granules, similarly to what we previously found in animals treated with the F-actin disrupting agents cytochalasin D and Latrunculin A, which completely block *de novo* F-actin assembly ^21,36,39^. This also suggests that the lattice provides a barrier to prevent or tune compound exocytosis once the signaling cascade eliciting exocytosis is triggered.

Interfering with either the Arp2/3-dependent machinery or the closely associated Ezrin significantly delays the integration of the granules without affecting their size or the F-actin lattice. Notably, impairment of the Arp2/3 function completely blocks the wave of branched actin polymerization; whereas, impairing Ezrin doesn’t affect the wave but prevents the formation of the F-actin ring, thus suggesting that the wave of branched actin is stabilized by Ezrin at the interface with the granule membrane. These data indicate that the Arp2/3-dependent step facilitates the integration but it is not essential. These actin filaments populations work in concert with the activity of NMIIA, as shown by the fact that: i) genetic ablation of NMIIA delays the integration to a similar extent to that of Arp2/3 disruption without affecting F-actin assembly ^39^, and ii) interference with Arp2/3 does not affect NMIIA recruitment (**Fig. 5**). How then does the Arp2/3 complex control granule integration? In several biological models It has been shown that arrays of branched actin form expanding gels that generate forces capable of pushing membranes ^71–74^. Therefore, it is plausible that the branched actin originating from and linked to the membranes by Ezrin expands toward the F-actin lattice and generates forces that are transmitted to the membranes. Alternatively, it is possible that the branched filaments/Ezrin module modulates a tension-driven membrane flow between the granules and the APM.

Both formins and the Arp2/3 complex have been shown to control different steps of exocytosis in different model systems. However, to our knowledge this is the first time that the strict coordination between these two modules is reported and, strikingly, in live animals under bona fide physiological conditions. We propose that it is very likely that these F-actin-based modules modulate the biophysical properties of granule membranes, including membrane tension, by exerting forces at a nanoscale level that could be transmitted via Ezrin to favor the exchange of membranes between granules and the APM. Although at the moment we cannot perform direct measurements of these membrane properties *in vivo*, based on our data we are confident that the integration of large granules *in vivo*, at least in the range of diameters we have described, does not rely on processes such as the folding of the granule membranes coupled with clathrin-mediated endocytosis (termed crumpling) ^75,76^.

In conclusion, these two coordinated force-generating modules that we describe in the salivary glands could potentially operate in other secretory systems where large micron-size vesicles, that release their contents in tubular compartments, are coated with an actomyosin complex after fusion (e.g., pancreas, lungs, lacrimal glands, endothelium) ^77–79^. However, since their role is to remodel membranes that are topologically and compositionally an extension of the plasma membrane it is likely that this machinery may play a role in other processes such as endocytosis or cell migration where membrane tension needs to be dynamically modulated.

## Supporting information

Heydecker et al. Supplementary Figures

Heydecker et al. Supplementary Table I

Heydecker et al. Supplementary Table II

Heydecker et al. Legends to Supplementary Movies

Movie S1 Heydecker et al.

Movie S2 Heydecker et al.

Movie S3 Heydecker et al.

Movie S4 Heydecker et al.

Movie S5 Heydecker et al.

Movie S6 Heydecker et al.

Movie S7 Heydecker et al.

Movie S8 Heydecker et al.

Movie S9 Heydecker et al.

## Acknowledgments

M.H. was supported by a Faculty of Medicine & Health Tuition Fee Scholarship and a Completion Scholarship from UNSW Sydney, and the Graduate Partnerships Program in the Office of Intramural Training & Education of the NIH.

R.W. was supported by the NIH, NCI, Center for Cancer Research Intramural Research Program (ZIA BC 011682).

K.N. was funded in whole or in part from the National Cancer Institute, National Institutes of Health, under Contract No. 75N91019D00024.

E.C.H. and P.W.G. were supported by grants from the Australian Research Council (DP160101623) and the Australian National Health and Medical Research Council (APP1100202, APP1079866).

We thank Dr. Renee Whan and members of the Katharina Gauss Light Microscope Facility at UNSW Sydney for support with live rodent imaging, and Dr. Maté Biro for his help with an earlier version of data quantification.

We thank Drs. Leonid Chernomordik (NICHD, NIH) and John Hammer III (NHLBI, NIH) for their critical reading of the manuscript and insightful comments.

The content of this publication does not necessarily reflect the views or policies of the Department of Health and Human Services, nor does mention of trade names, commercial products, or organizations imply endorsement by the U.S. Government.

## Author Contributions

M.H. designed and performed the experiments, analyzed the data, and wrote the manuscript.

A.S., M.T., A.M., S.E. performed experiments with inhibitors.

M.A.A. performed *in vivo* transfections.

K.N., A.B., D.T. performed and analyzed FIB-SEM data.

J.L.G.N performed the qPCR and analysis.

D.C. performed quantitative analysis.

E.C.H. conceived of and supervised the project, edited the manuscript, and acquired funding.

P.W.G. conceived of and supervised the project, edited the manuscript, and acquired funding.

R.W. conceived of and supervised the project and wrote the manuscript.

## Declaration of Interests

P.W. Gunning and E.C. Hardeman are Directors and shareholders of TroBio Therapeutics Pty Ltd, a company that is commercializing anti-tropomyosin drugs for the treatment of cancer. Their labs receive funding from TroBio to evaluate anti-tropomyosin drug candidates. All authors declare no financial competing interests. All authors declare no non-financial competing interests.

## Methods

### Animal strains and procedures

All experiments using animals were performed in accordance with the guidelines provided by: 1) the National Cancer Institute (National Institutes of Health, Bethesda, MD, USA) Animal Care and Use Committee and were compliant with all relevant ethical regulations regarding animal research; and 2) the NSW Animal Research Act (1985) and Australian National Health and Medical Research Council (NHMRC) ‘Code’ 8th edition (2013). All experiments were approved by the UNSW Sydney Animal Care and Ethics Committee under application 17/100B. Male or female mice (age 8-24 weeks) and male and female rats (age 6-10 weeks) were used in this study. Wistar/Sprague Dawley rats (150 g) were purchased from the Animal Resources Centre, Perth, WA, Australia. Standard laboratory chow and water was provided ad libitum. The mTom/mGFP Cre reporter mouse ^49^ was purchased from Jackson Laboratory. Hemizygous Lifeact-GFP and Lifeact-RFP transgenic mice were a gift from Roland Wedlich-Soldner ^80^. GFP-NMIIA knock-in (KI) mice were generated as described ^81^ and crossed with the RFP-Lifeact strain. The mouse line expressing Tpm3.1 C-terminally tagged with mNeonGreen (NG) at the endogenous locus [B6-Tpm3tm5(mNeonGreen)Hrd] was generated as described ^82^. Conditional mDia1 (mDia1^fl/fl^) ^54^ or ARPC2 (ARPC2^fl/fl^) ^55^ flox mice were crossed with mice harboring (TMX)-inducible Cre recombinase under the control of the Mist1 promoter ^56^ and the mT/mGFP reporter ^49^ (**Fig. S3A-C**; _mDia1_fl/fl Mist1-Cre, mT/mG_, ARPC2_fl/fl Mist1-Cre, mT/mG_). Cre recombinase was induced by_ two intraperitoneal injections of Tamoxifen (75 mg/kg body weight), two days apart. Mice were imaged 3-4 weeks after the first injection. Cre-expressing cells (GFP positive) and non Cre-expressing cells (mTom) could be differentiated by their fluorescence and their secretory granules visualized after their fusion with the APM39,49

### Intravital subcellular microscopy

Mice (18-34 g) were anesthetized by i.p. injection of 100 mg/kg ketamine/15 mg/kg xylazine. Salivary glands were surgically externalized and prepared for intravital subcellular microscopy as described ^83^. Salivary glands were stabilized on a coverslip on the microscope stage with vasculature and innervation functionally intact.

#### Instruments

For high resolution 3D imaging a Nikon Ti2 spinning disk microscope was utilized. See Supplementary Table 1 for details of the spinning-disc head, camera and objectives used. For live imaging, the acquisition speed was set at 80–300 ms per frame, with Z-stacks acquired at 0.05–0.3 μm apart. Nikon Elements Software was used for post-acquisition image alignment to correct for image drift, denoising and for 3D deconvolution using the ‘Automatic’ option, which used the Richardson– Lucy algorithm ^84^ with: i) an additional step for pre-noise estimation, and ii) automatic determination of the number of iterations. For image acquisition from fixed tissue sections, the sampling frequency at the camera sensor was 45 nm which was enabled by a x1.5 tube lens. Although this frequency was well above the required Niquist criterion, the oversampling allowed for an extended dynamic range (when combined with the 16-bit analog-to-digital converter and longer exposure times) as has been used previously ^85^, and, together with 3D deconvolution, allowed an improvement in highlighting high-frequency features in the images ^86^.

Intravital imaging was also performed using a Nikon A1 inverted laser scanning confocal microscope fitted with a CFI Plan Apochromat lambda series 60x/1.27NA water immersion objective, an Okolab humidified temperature-controlled microscope enclosure, objective heater and a custom-made stage insert. Animal body and salivary gland temperatures were monitored with a MicroTherma T2 thermometer equipped with rectal and thin implantable probes (Braintree Scientific) and heating adjusted to maintain 37^0^C body and SG temperature. GFP and mNG were excited with a 488 nm laser, and RFP and tdTomato were excited with a 561 nm laser. For time-lapse imaging, frames were acquired in sequential mode at 2.019 sec/frame or 241 msec/frame at spatial sampling of 100 nm per pixel. An imaging plane with a visible canaliculus or apical membrane was selected as close to the surface of the organ as possible – at depth of 10-20 μm for rat and 8-12 μm for mouse salivary glands. During imaging, any noticeable drift was manually corrected in X, Y or Z dimension. For some experiments, drugs (200 μM CK666 or 200 μM SMIFH2, Sigma-Aldrich Pty Ltd; 250 μM NSC668394, EMD Millipore) or 1.5% DMSO solutions were prepared using pre-warmed 37^0^C saline and perfused over the surface of the gland at 10 μL/min during imaging using a PHD Ultra Nanomite programmable syringe pump (Harvard Apparatus). Exocytosis of secretory granules was stimulated by subcutaneous injection of isoproterenol (ISOP) at 0.025 mg/kg (rats) or 0.01 mg/kg (mice).

### Image acquisition and data processing

Images were recorded using NIS Elements software and either a Nikon A1 confocal or Nikon Ti2 spinning disk microscope. Image processing and data extraction was performed using Imagej/Fiji ^87^ NIS Elements and Imaris. Drift and motion correction on time lapse image stacks was carried out with the StackReg ImageJ plugin ^88^. Nikon Elements Software was used for post-acquisition image alignment and depending on the signal to noise ratio the Nikon Denoise.ai function was used followed by 3D deconvolution using the Richardson–Lucy algorithm. The Richardson–Lucy/maximum likelihood image restoration algorithm was used for fluorescence microscopy with: i) an additional step for pre-noise estimation, and ii) automatic determination of the number of iterations.

#### Diameters of granules and F-actin thickness

Diameters of granules were measured using the line scan function in Fiji. At first detection of mTom/mGFP a line scan was drawn across the center of the maximum diameter. At each end of the membrane, the fluorescence displays a Gaussian intensity profile and the peak-to-peak value was measured as the diameter. For actin diameters the same strategy was used; during thickening the intensity formed plateaus instead of displaying distinct peaks. The length of intensity plateaus was measured as diameter for thickening actin. Diameters were then exported to Excel and their average and SD calculated. The data was plotted using GraphPad Prism and the first significant decrease in diameter values was determined using the one-way ANOVA test (package in Prism).

For actin thickness, a line scan was drawn across the maximum diameter and the plateau on each side was measured, added and divided by 2. The actin diameter was plotted against the actin thickness and linear regression performed using Prism.

#### Surface area

Surface area (SA) was calculated by approximating the granule to a sphere and using the formula SA=4*π*(diam/2)^2^. The rate of surface area change was calculated as the difference in SA between two time points in the time-lapse images divided by the time interval. Statistical analysis for the initial diameter (**Fig. 4C,E**) was performed using the one-way ANOVA test (package in Prism).

Granule exocytosis events from 3-8 animals were quantified, as specified in figure legends. Images for the figures were adjusted for contrast and brightness in Imaris or ImageJ to have an optimal display range for features of interest, such as secretory granules and then converted to RGB images.

### Cardiac fixation and cryosectioning of the salivary gland

For cardiac perfusion, the left ventricle of the heart was punctured in anesthetized mice, and PBS injected to remove blood from the right atrium, followed by 25 mL fixative (4% formaldehyde in PBS, pH 7.3). Glands were then excised and placed in fixative for 30 min at RT. After fixation, salivary glands were put through a sucrose gradient (10% → 20% → 30%→ 30%:OCT [1:1]) for cryoprotection. Once the tissue sank in the 30% sucrose:OCT [1:1] solution, it was placed in a mold in OCT and frozen on dry ice. Cryosections (10 μm) were cut and adhered to Superfrost Plus slides (Electron Microscopy Sciences).

### Indirect Immunofluorescence

Cryosections were washed 3x5 min in PBS, permeabilized in 0.5% Triton X-100 in PBS for 20 min, and blocked in 10% NGS in PBS for 1 hr at RT. Sections were then incubated with the respective primary antibody (see **Supplementary Table 1**) overnight at 4^0^C. Sections were rinsed 3x with PBS and stained with respective secondary antibodies (see **Supplementary Table1**). Sections were then rinsed 3x with PBS and mounted using Fluormount-G TM (Invitrogen) on a glass slide with no. 1.5 coverslips.

### Inhibitor treatment of salivary glands in live mice

Mouse salivary glands were surgically exposed and bathed in 200 μM CK666, 200 μM SMIFH2 or 250 μM NSC in saline at 37^0^C, 10 min. Dimethylsulfoxide (1.5%) was used as a control. Mice were then injected subcutaneously with 0.01 mg/kg ISOP to stimulate exocytosis. After 30 min, glands were covered with Parafilm and cardiac fixation was performed as described above.

### Plasmid DNA preparation

DNA was extracted from bacterial overnight cultures using Qiagen EndoFree Plasmid Kit (Qiagen). Plasmid DNA was stored in TE-Buffer and diluted to 1 μg/μl aliquots. Plasmid constructs mDia1-Emerald (Addgene plasmid # 54157) and Arp2-Emerald (Addgene plasmid # 53992) were a gift from Michael Davidson. Lifeact-RFP construct was a gift from Roland Wedlich-Soldner ^80^.

### *In vivo* transfections in rats

Rats (150–225 g) were obtained from the Animal Resources Centre, Perth, WA, Australia and allowed to acclimate for 1 week. Rats were anesthetized and salivary glands transfected as previously described ^89^ with the following modifications: 24 μg of plasmid DNA was mixed with Lipofectamine™ 3000 according to the manufacturer’s instructions.

### Real-time quantitative reverse transcription PCR (qRT-PCR) of gene expression

Submandibular glands were isolated and dissociated with 1 mg/ml collagenase IV (Sigma-Aldrich) for 20 min at 37^0^C (shaking at 800 rpm). Samples were filtered through 70 μm cell strainers to obtain single cell suspensions. The acinar glands were resuspended in a final volume of 2 mL FACS wash buffer (2% HI-FCS, 2 mM EDTA, 0.02% sodium azide in 1x PBS). A total of 1x10^6^ GFP-expressing acinar cells and non-expressing cells were sorted using BD InfluxTM flow sorter (BD Biosciences; Franklin Lakes, NJ, USA). Sorted cells were then centrifuged (300xg, 5 min) before RNA isolation using the RNeasy Mini Kit (Qiagen), according to manufacturer’s instructions. 1.5 μg of RNA was reverse transcribed to cDNA using the High-Capacity cDNA Reverse Transcription Kit (Applied Biosystems, ThermoFisher Scientific) according to manufacturer’s instructions. qRT-PCR was carried out using primers specific for each actin nucleator. The primers were predesigned and synthesized by Sigma-Aldrich (KiCqStart® SYBR® Green Primers). 20 ng cDNA was added to each well of a 96-well PCR plate with 1 μM forward and reverse primer for each nucleator. 40 cycles were performed, with denaturing temperature at 95^0^C for 15 sec, annealing at 55^0^C for 30 sec, and extension at 72^0^C for 30 sec. The amount of the PCR product was determined in real time by using SYBR Green and detected in a BIO-RAD CFX96 real-time system (Bio-Rad Laboratories, Hercules, CA, USA). The amount of gene expression was calculated by using the ΔΔCT (cycle threshold) method, where ΔCT = CT (gene of interest) – CT (housekeeping gene) describes the difference in CT values and ΔΔCT = ΔCT (treated sample) – ΔCT (untreated sample) describes the difference between the control and the experimental sample. The data was normalized to the expression of the housekeeping genes β2-microglobulin (β2M) and ribosomal protein L13A (RPL13A) and with a universal positive control. Melting curve analysis was done for confirmation of product specificity (list of primers in Supplementary Table 2).

### FIB SEM

#### Sample processing

Salivary glands were fixed in in 2% glutaraldehyde for 2 hr at RT, followed by washes in 0.1M sodium cacodylate buffer. Glands were stained in 1% osmium tetroxide, washed again in cacodylate buffer, distilled water and then stained in 0.5% uranyl acetate in 0.05 M sodium maleate buffer. Glands were dehydrated through an ethanol gradient and embedded in PELCO Eponate 12^TM^. Once sample blocks were cured, the block surface was milled flat using an ultramicrotome to expose the glands on the block surface. Sample blocks were mounted on imaging stubs using conductive glue and prepared for microscope loading.

#### Sample acquisition

FIB-SEM images of salivary acini were typically acquired at 5 nm pixel sampling and 15 nm FIB mill step size, and the resulting stacks of 8-bit .tiff images were aligned, inverted and binned to generate .mrc volumetric reconstructions at 15 nm cubic voxel sampling, using established protocols ^90^. Sub-volumes containing canaliculi were then excised computationally and segmented out using 3D Slicer (www.slicer.org); a combination of threshold-based segmentation and manual clean-up captured both the ducts as well as fusing salivary granules. Visual inspection revealed several fusing granules that were then excised at the necks to yield label maps of every instance of fusing granules captured in a given volume EM reconstruction. These were converted into 3D meshes for downstream calculations.

#### Volumetric measurements

The 3D Slicer quantification module was used to calculate the volume (V), surface area (SA) and roundness (deviation from sphericity; perfect sphere = roundness of 1) of each fusing vesicle. Granule diameters were calculated mathematically from V and SA assuming sphericity; these were cross checked with manual representative measurements at the widest points of the granule cross-sections parallel to the axis of the canaliculus. The measurements were very close except for the smallest vesicles, where there was significant deviation from sphericity, so manual measurements were used throughout.

To test the curvature of these granules, we exported the meshes and used the Trimesh library in Python (https://pypi.org/project/trimesh/) to generate discrete mean curvature measures of each granule. This function returns a mean curvature calculated from the dihedral angles between all planes emanating from each vertex of the mesh. An LUT was applied to the mean curvature value at each vertex from +1 (convex, red) through 0 (planar, white) to -1 (concave, blue) and mapped on to the original mesh vertices. This shows visually where and how each granule is curved. The mean of all the mean curvatures per granule was calculated and plotted.

