## Supplementary material for "Spatial and Temporal Coordination of Force-generating Actin-based Modules Drives Membrane Remodeling *In Vivo*": Heydecker et al. Supplementary Figures

### Supplementary Information

### Figure S1

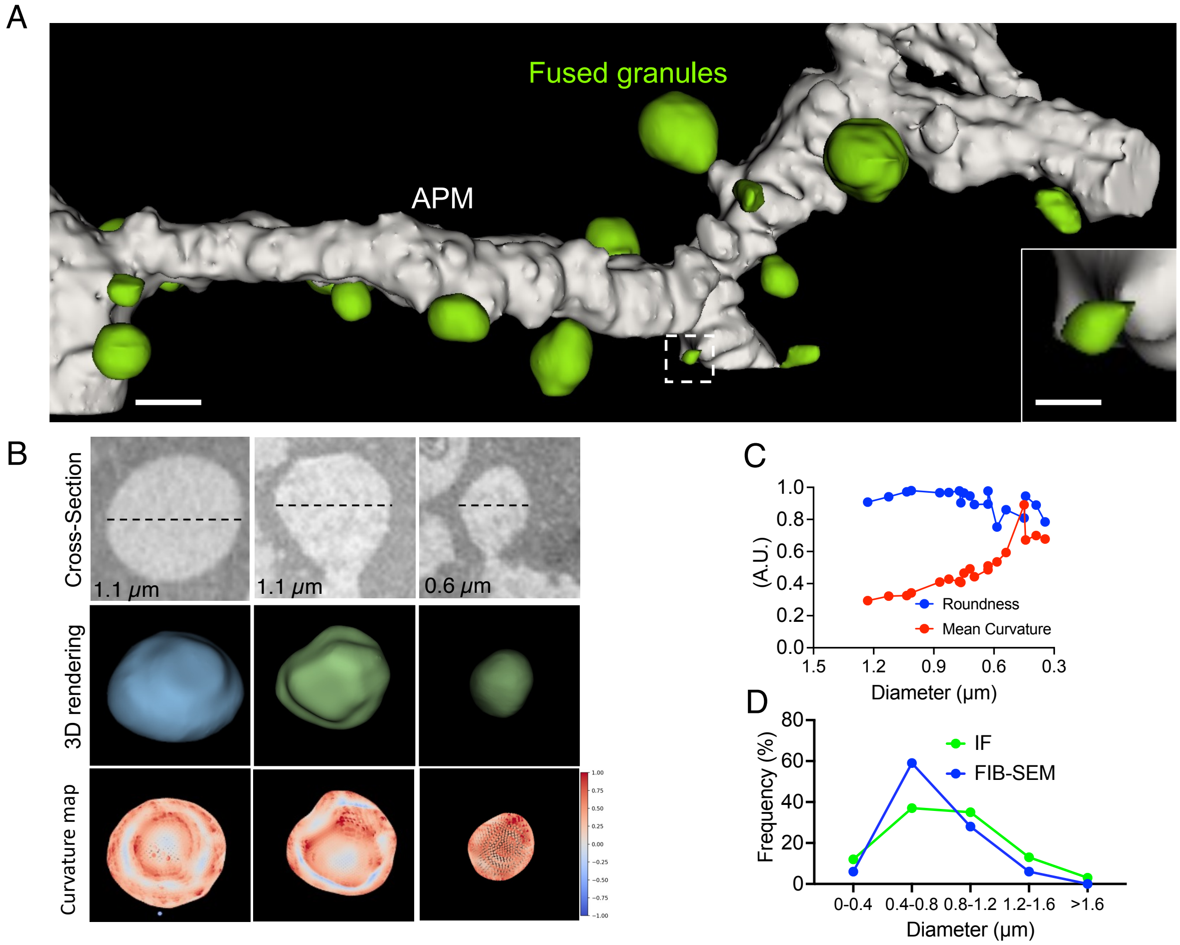

**Figure S1. FIB-SEM of salivary gland secretory granules**. WT mice were injected with 0.01 mg/kg ISOP and after 10 min the glands were processed for FIB-SEM and imaged. **A.** Volume rendering of a segment of the APM (white) and the fused secretory granules (green). Bar 1 µm. Inset shows a small vesicle of undetermined origin. Bar 200 nm. **B**. Example of secretory granules: non-fused (left) and at two different stages of integration (center and right). Upper panels - EM micrographs. Center panel - Volume rendering. Lower panels - Heat map of local curvature (red positive, blue negative). **C.** Analysis of roundness and mean curvature of fused secretory granules. N=19 granules in 2 animals. **D**. Distribution of the granule diameters measured by FIB-SEM (N=19 granules in 2 animals) or IVM (N=55 granules in 5 animals).

### Figure S2

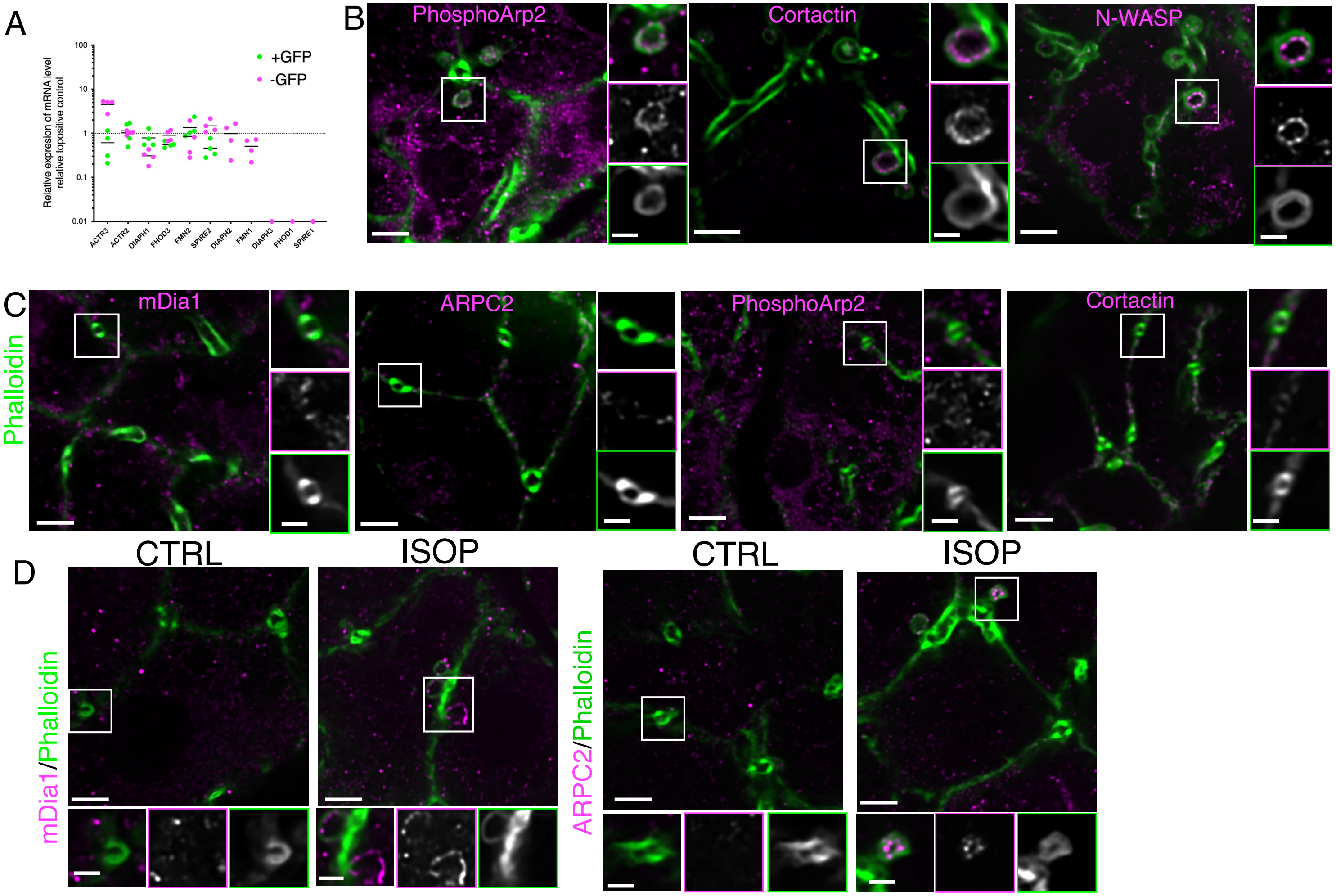
A
Relative expresion of mRNA level relative topositive control
100
10
1
0.1
0.01
+GFP
-GFP
ACTR3
ACTR2
DIAPH1
FHOD3
FMN2
SPIRE2
DIAPH2
FMN1
DIAPH3
FHOD1
SPIRE1
B
PhosphoArp2
Cortactin
N-WASP
C
Phalloidin
mDia1
ARPC2
PhosphoArp2
Cortactin
D
CTRL
ISOP
mDia1/Phalloidin
ARPC2/Phalloidin

**Figure S2**. **Identification of the actin nucleators in mouse and rat salivary glands**. **A**. Acinar cells were examined for actin nucleator mRNA expression. Salivary glands of the acinar-specific Mist1-inducible/membrane reporter mTom/mGFP mouse were transfected with an AAV9 Cre-vector by injection into the Wharton duct. Cells were sorted by their fluorescence (+GFP, -GFP) to identify acinar cells (+GFP) and actin nucleator transcript levels were measured using qPCR. **B,C**. WT mice were either injected with 0.01 mg/kg ISOP (**B**) or left untreated (**C**). Salivary glands were processed for cryosectioning and indirect immunofluorescence, and labeled with Alexa 647 phalloidin (green) and antibodies raised against phospho-Arp2, ARPC2, N-WASP or cortactin (Alexa Fluor-488 goat anti-rabbit, magenta). Insets show individual granules (**B**) or the APM (**C**). Bar 3 μm; 1 μm insets. **D**. Wistar rats were either injected with 0.025 mg/kg ISOP or left untreated (CTRL). Salivary glands were processed for cryosectioning and indirect immunofluorescence, and labeled with Alexa 647 phalloidin (green) and antibodies raised against mDia1 or ARPC2 (Alexa Fluor-488 goat anti-rabbit, magenta). Insets show individual granules (ISOP or the APM (CTRL). Bar 3 μm; 1 μm insets.

**Figure S3**

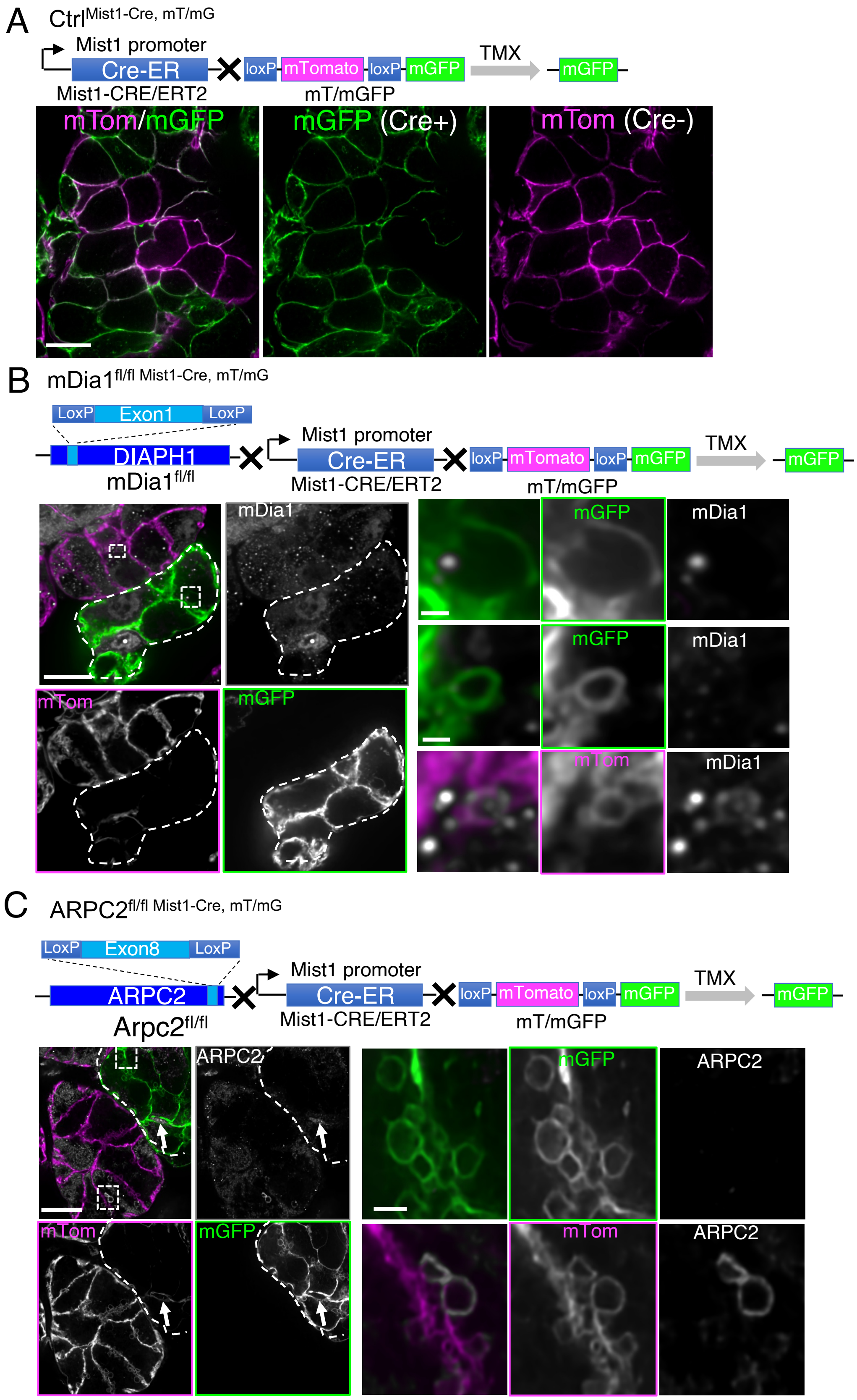

A
Ctrl Mist1-Cre, mT/mG
Mist1 promoter
Cre-ER
loxP
mTomato
loxP
mGFP
TMX
mGFP
Mist1-CRE/ERT2
mT/mGFP
mTom/mGFP
mGFP (Cre+)
mTom (Cre-)
B
mDia1 fl/fl Mist1-Cre, mT/mG
LoxP
Exon1
LoxP
DIAPH1
mDia1 fl/fl
Mist1 promoter
Cre-ER
Mist1-CRE/ERT2
loxP
mTomato
loxP
mGFP
mT/mGFP
TMX
mGFP
mDia1
mGFP
mDia1
mGFP
mDia1
mTom
mGFP
mTom
mDia1
C
ARPC2 fl/fl Mist1-Cre, mT/mG
LoxP
Exon8
LoxP
ARPC2
Arpc2 fl/fl
Mist1 promoter
Cre-ER
Mist1-CRE/ERT2
loxP
mTomato
loxP
mGFP
mT/mGFP
TMX
mGFP
ARPC2
mGFP
ARPC2
mTom
mGFP
mTom
ARPC2

**Figure S3**. **Tamoxifen-inducible knock down of mDia1 and ARPC2**. **A.** Mouse expressing Cre recombinase under the acinar-specific Mist1 TMX-inducible promoter and the membrane reporter mTom/mGFP ($Ctrl^{Mist1\text{-}Cre,\ mT/mG}$). Mice were treated with 75 mg/kg TMX for 4 weeks and salivary glands exposed and imaged by confocal microscopy. Acinar cells positive for GFP express Cre recombinase. Bar 10 µm. **B**. Mice in which exon 1 of the gene encoding mDia1 (DIAPH1) is flanked by LoxP sites were crossed with the acinar-specific Mist1 TMX-inducible promoter/membrane reporter mTom/mGFP mice ($mDia1^{fl/fl\ Mist1\text{-}Cre,\ mT/mG}$). After 4 weeks of TMX treatment, salivary glands were exposed, processed for indirect immunofluorescence, and labeled with antibodies directed against mDia1. **C.** Mice in which exon 8 of the gene encoding Arpc2 (ARPC2) is flanked by LoxP sites were crossed with the acinar-specific Mist1 TMX-inducible promoter/membrane reporter mTom/mGFP mice ($ARPC2^{fl/fl\ Mist1\text{-}Cre,\ mT/mG\ mG}$). After 4 weeks of TMX treatment, mice were injected with 0.01 mg/kg ISOP and salivary glands excised, processed for indirect immunofluorescence, and labeled with antibodies directed against ARPC2. Cells that express Cre recombinase (green and highlighted by dashed lines) have lower levels of ARPC2 than those that do not (magenta). Bar 10 µm. Insets show that in cells that do not express Cre (green), ARPC2 is not present on the fused granules. Bar 1 µm.

**Figure S4**

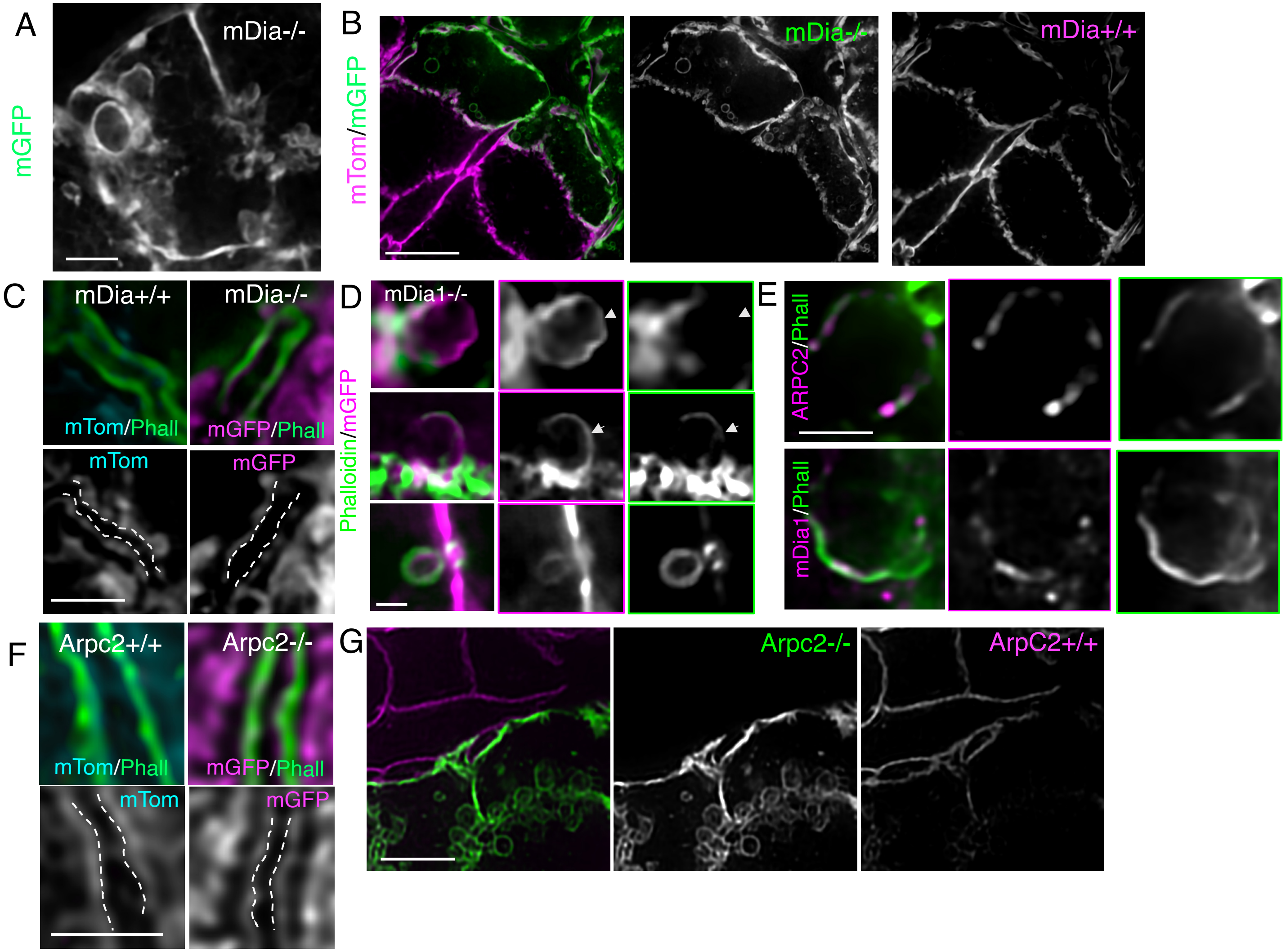

A
mDia-/-
mGFP
B
mTom/mGFP
mDia-/-
mDia+/+
C
mDia+/+
mDia-/-
mTom/Phall
mGFP/Phall
mTom
mGFP
D
mDia1-/-
Phalloidin/mGFP
E
ARPC2/Phall
mDia1/Phall
F
Arpc2+/+
Arpc2-/-
mTom/Phall
mGFP/Phall
mTom
mGFP
G
Arpc2-/-
ArpC2+/+

**Figure S4**. **Effect of mDia1 and ARPC2 depletion/inhibition in mouse salivary glands**. mDia1$^{fl/fl\ Mist1\text{-}Cre,\ mT/mG}$ (**A-D**) and ARPC2$^{Mist1\text{-}Cre,\ mT/mG}$ (**F,G**) mice were injected with 75 mg/kg of TMX. After 4 weeks they were injected with 0.01 mg/kg of ISOP (**B-D,F,G**) and after 10 min glands were processed for indirect immunofluorescence and imaged with spinning disk microscopy. Some mice were imaged after 6 weeks of TMX injection without ISOP stimulation (**A**). **A.** Accumulation of endomembranes in mDia-/- cells (expressing GFP). Bar 3 µm. **B**. Enlarged granules accumulated in mDia -/- but not in mDia+/+ cells. Bar 5 µm. **C,D.** Salivary glands were labelled with phalloidin (green). **C**. mDia1 depletion did not affect the diameter of the APM. Bar 1 µm. **D**. Enlarged granules did not recruit F-actin or in reduced amounts (arrowheads, upper and center panels), when compared with normal size granules (lower panel). Bar 1 µm. **E**. WT mice were treated with 200 µM SMIFH2. Salivary glands were excised, processed for indirect immunofluorescence and labeled with phalloidin (green), and antibodies against mDia1 (magenta, lower panel) or ARPC2 (magenta, upper panel). Bar 2 µm. **F**. ARPC2 depletion does not affect the diameter of the APM. **G**. Accumulation of normal size granules only in ARPC2 -/- but not in ARPC2+/+ cells. Bar 4 µm.

**Figure S5**

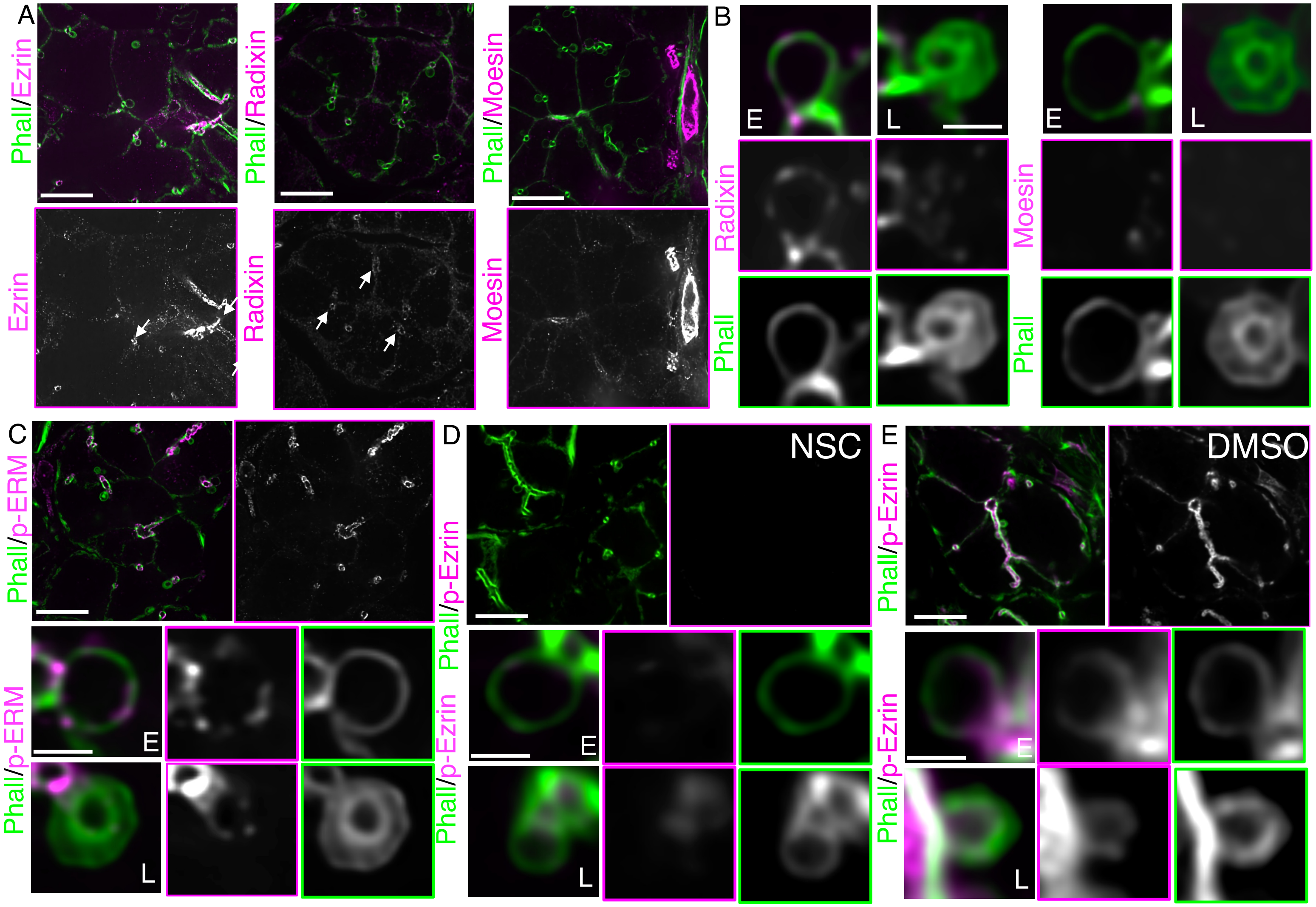

A
Phall/Ezrin
Phall/Radixin
Phall/Moesin
Ezrin
Radixin
Moesin
B
E
L
E
L
Radixin
Moesin
Phall
Phall
C
Phall/p-ERM
E
L
D
NSC
Phall/p-Ezrin
E
L
E
DMSO
Phall/p-Ezrin
E
L

**Figure S5. Expression, localization and function of ERM proteins**. **A-C.** WT mice were left untreated (**A-C**) or treated for 30 min with 250 µM NSC (**D**) or vehicle (DMSO, **E)**. Mice were injected with 0.01 mg/kg ISOP and after 10 min salivary glands were excised and processed for indirect immunofluorescence with Alexa488-phalloidin (green) and antibodies directed against Ezrin (**A**), Radixin (**A,B**), Moesin (**A,B**), pan phospho-ERM (**C**) or phosphor-Exrin (**D,E**). **A**. Low mag of acini showing the localization of the various markers (magenta, arrows) at the APM. Only Radixin and Ezrin localize at the APM (arrows). Bar 10 µm. **B**. Radixin localizes on the secretory granules at early (E) but not late (L) stages (arrowheads). Moesin is not recruited on the fused granules. Bar 1 µm. **C**. pERM is detected on the secretory granules at both early (E) and late (L) stages (arrowheads). Bar 10 µm. **D,E**. Treatment with NSC reduces the levels of p-Ezrin phosphorylation. Bar 10 µm; 1 µm (high magnification).
