## Supplementary material for "Spatial and Temporal Coordination of Force-generating Actin-based Modules Drives Membrane Remodeling *In Vivo*": Heydecker et al. Supplementary Table I

| Primary | Source | Cat. # | Dilution | Figure |
|---|---|---|---|---|
| Alexa 488 Phalloidin | Invitrogen | A12379 | 1:250 | Fig.1E-Fig.2A,B-Fig.3B,E-Fig.4B-Fig.5G-F-Fig.S5D,E |
| Alexa 405 Phalloidin | Invitrogen | A30104 | 1:250 | Fig.5D-Fig.6H-Fig.S4C,F |
| Alexa 568 Phalloidin | Invitrogen | A12380 | 1:500 | Fig. 4D-Fig.6A-D,G,H-Fig.S2C,D-Fig.S6A-D |
| Alexa 647 Phalloidin | Invitrogen | A-22287 | 1:200 | Fig.S2B |
| Anti-DIAPH1 (mDia1) | G-biosciences | ITA2946 | 1:200 | Fig.2A-Fig.5G Fig.S2C,D-Fig. S3B-Fig.S4E |
| Anti-Arpc2 | Sigma | HPA008352 | 1:500 | Fig.2A-Fig.5D,G-Fig.S2C,D-Fig. S3C-Fig.S4E |
| Anti Tpm3.1 anti-γ/9d | | | 1:150 | Fig.3A,B |
| Anti-Non-muscle Myosin Heavy Chain IIA | Biolegend | 909802 | 1:500 | Fig.3C,3E-Fig.4G-Fig.5D,G |
| Anti-Ezrin | Invitrogen | MA5-13862 | 1:500 | Fig.6C,D-Fig.S5A,B |
| Anti phospho-Arp2 (Thr237, Thr238) | Invitrogen | PA5-143650 | 1:500 | Fig.S2B,C |
| Anit-Cortactin | Cell Signaling Technologies | 3503S | 1:500 | Fig.S2B,C |
| Anti N-Wasp | Cell Signaling Technologies | 4848S | 1:500 | Fig.S2B,C |
| Anti-Phospho-Ezrin (Thr567) | Invitrogen | PA5-37763 | 1:500 | Fig.6G-Fig-S5K,L |
| Anti-Moesin | Abcam | ab52490 | 1:500 | Fig.S5E,F |
| Anti-Radixin | Cell Signaling Technologies | 2636S | 1:500 | Fig.S5C,D |
| Anti-phospho ERM | Cell Signaling Technologies | 3141S | 1:500 | Fig.S5G,H |
| **Secondary** | | | | |
| Alexa Fluor-488 Goat anti-Rabbit | Invitrogen | A-11070 | 1:500 | Fig.3A,B,D,E-Fig.6A-D,G,H- Fig.S2B,D-Fig.S6A-C- |
| Alexa Fluor-568 Goat anti-Rabbit | Invitrogen | A-21430 | 1:500 | Fig.4G-Fig.5G-FigS5A-C |
| Alexa Fluor-647 Goat anti-Rabbit | Invitrogen | A-21244 | 1:500 | Fig.5D-Fig.S5D,E |
