## Supplementary material for "Spatial and Temporal Coordination of Force-generating Actin-based Modules Drives Membrane Remodeling *In Vivo*": Heydecker et al. Supplementary Table II

| Gene | Forward | Reverse |
|---|---|---|
| Actr2 | GTTGAGTCATATACACTTCCAG | CTGAAATAAAGCCTCTGGTG |
| Actr3 | CATCCAGAGTTTGCTAATCC | TGGCATCTACAGTTCTCTTC |
| Diap1 | CTGGACGGATTAAAAAGTGG | TTTTCAATCTCTCGAAGCTC |
| Diap2 | GAAAGGAAATGGAAGAGAAGAG | CTCATCACCCTCTTTGTTTATG |
| Diap3 | GGAAAGCATCCTTGAAGAAG | ATCAGGAGATGTAACGAGAG |
| Fhod1 | CAATGCTATCTTGGAAAAGC | CTTGTCTTCCTGGAATATCTG |
| Fhod3 | TGCTGGAGCAGTTTAATATC | TTCTGGATTGAGTGTCTACG |
| Fmn1 | CTTATATGAAAATCGAGCCCAG | TGTAAGGATGTGATACCCTC |
| Fmn2 | TAGTAAAAGAGACTCCAGCC | GAATTCCTACTGCTTGTGAC |
| Spire1 | GGAAGAAGCCTCCAAAATTC | TAGCAGAATTCCTCAAGAGAC |
| Spire2 | AAGAGGGGGAAGATTTGC | CTTGGAAGGCATCTTCATC |
| β2M | GTATGCTATCCAGAAAACCC | CTGAAGGACATATCTGACATC |
| Rpl13a | CCTATGACAAGAAAAAGCGG | CAGGTAAGCAAACTTTCTGG |
