## Supplementary material for "Spatial and Temporal Coordination of Force-generating Actin-based Modules Drives Membrane Remodeling *In Vivo*": Heydecker et al. Legends to Supplementary Movies

**Movie S1. Time-lapse of secretory granule integration in a mouse expressing GFP-LF and mTom.** GFP-LF (green) is recruited on the granule (mTom, magenta), and as the integration initiates the external diameter of the lattice does not significantly change. Polymerization of actin continues and manifests as an increase in the thickness of GFP-LF around the granules (still images in **Fig. 1C**). Single focal plane acquired by spinning disk confocal microscopy. This experiment was repeated in 5 mice.

**Movie S2. Time-lapse of secretory granule integration in a rat transiently transfected with RFP-LF and mDia1-Emerald.** A first pool of mDia1-emerald (magenta) appears on the granules ~2-3 sec before the RFP-LF (green) followed by a second pool later (still images in **Fig. 2C**). Single focal plane acquired by confocal microscopy. This experiment was repeated in 2 rats.

**Movie S3. Time-lapse of secretory granule integration in a rat transiently transfected with RFP-LF and Arp2-Emerald.** Arp2-Emerald (magenta) is recruited on the granule after the appearance of RFP-LF (green) and appears to localize beneath the lattice (still images in **Fig. 2D**). Single focal plane acquired by confocal microscopy. This experiment was repeated in 2 rats.

**Movie S4. Time-lapse of secretory granule integration in a mouse expressing Tpm3.1-NG and RFP-LF.** Tpm3.1-NG (magenta) is recruited on the granules with the RFP-LF (green) and associates with the lattice during the integration (still images in **Fig. 3D**). Single focal plane acquired by confocal microscopy. This experiment was repeated in 3 mice.

**Movie S5. Time-lapse of secretory granule integration in a mouse expressing GFP-NMIIA and RFP-LF.** GFP-NMIIA (magenta) is recruited on the granules after RFP-LF (green) and associates with the lattice during the integration (still images in **Fig. 3F**). Single focal plane acquired by confocal microscopy. This experiment was repeated in 3 mice.

**Movie S6. Time-lapse of secretory granule integration in mDia1 floxed mice.** Expansion of a secretory granule labeled with the membrane marker mGFP in a mouse depleted of mDia1 (left panel). Right panel shows granule integration in a control mouse. Maximal projections of 3D stacks acquired in time-lapse (still images in **Fig. 4B**). This experiment was repeated in 8 (mDia-/-) and 5 (mDia+/+) mice.

**Movie S7. Time-lapse of secretory granule integration in ARPC2 floxed mice.** Delayed integration of a secretory granule labeled with the membrane marker mGFP in a mouse depleted of ARPC2 (left panel). Right panel shows granule integration in a control mouse. Maximal projections of 3D stacks acquired in time-lapse (still images in **Fig. 5B**). This experiment was repeated in 4 (APRC2-/-) and 5 (APRC2+/+) mice.

**Movie S8. Time-lapse of secretory granule integration in GFP-LF/mTom mice treated with CK666.** The integration of a secretory granule (mTom, magenta) in a mouse treated with CK666 is significantly delayed and occurs without GFP-LF thickening (green) (still images in **Fig. 5F**). This experiment was repeated in 5 mice.

**Movie S9. Time-lapse of secretory granule integration in GFP-LF/mTom mice treated with NSC668394.** The integration of a secretory granule (mTom, magenta) in a mouse treated with NSC668394 is significantly delayed and occurs without the formation of GFP-LF (green) ring (still images in **Fig. 6E**). Single focal plane acquired by spinning disk confocal microscopy. This experiment was repeated in 3 mice.
